# Selective modification of ascending spinal outputs in acute and neuropathic pain states

**DOI:** 10.1101/2024.04.08.588581

**Authors:** David A. Yarmolinsky, Xiangsunze Zeng, Natalie MacKinnon-Booth, Caitlin Greene, Chloe Kim, Clifford J. Woolf

**Affiliations:** Boston Children’s Hospital, F.M. Kirby Neurobiology Center, Boston, MA, United States; Department of Neurobiology, Harvard Medical School, Boston, MA, United States

## Abstract

Pain hypersensitivity arises from the plasticity of peripheral and spinal somatosensory neurons, which modifies nociceptive input to the brain and alters pain perception. We utilized chronic calcium imaging of spinal dorsal horn neurons to determine how the representation of somatosensory stimuli in the anterolateral tract, the principal pathway transmitting nociceptive signals to the brain, changes between distinct pain states. In healthy conditions, we identify stable, narrowly tuned outputs selective for cooling or warming, and a neuronal ensemble activated by intense/noxious thermal and mechanical stimuli. Induction of an acute peripheral sensitization with capsaicin selectively and transiently retunes nociceptive output neurons to encode low-intensity stimuli. In contrast, peripheral nerve injury-induced neuropathic pain results in a persistent suppression of innocuous spinal outputs coupled with activation of a normally silent population of high-threshold neurons. These results demonstrate the differential modulation of specific spinal outputs to the brain during nociceptive and neuropathic pain states.

## INTRODUCTION

A fundamental question in neurobiology is how the same fixed external stimulus may evoke different perceptual, emotional, and behavioral responses as a consequence of changes in internal state^1–3^. An important example of this phenomenon is the transition between protective nociceptive pain, in which the sensation of pain is triggered only by an imminent danger of tissue damage by a noxious stimulus, to a sensitized pain state, in which intense pain is now elicited by innocuous stimuli^4^. Acute pain hypersensitivity following tissue injury or inflammation, drives adaptive behaviors that temporarily protect the healing tissues from further harm^5,6^. Conversely, in chronic conditions, such as neuropathic pain, a persistent and inappropriate perception of pain maladaptively impairs motor, cognitive, and emotional functions without providing any protective benefit^7^.

The induction of pain hypersensitivity is initiated by plasticity within the first two stations of somatosensory processing, in primary sensory and spinal dorsal horn neurons. Both undergo functional alterations in response to intense or damaging stimuli, respectively referred to as peripheral and central sensitization. The functional plasticity of primary sensory neurons is mediated by diverse mechanisms, including changes in the transduction sensitivity of nociceptor peripheral terminals due to post-translational modification and membrane trafficking of ion channels, bi-directional neuro-immune interactions and altered gene expression^8–10^, while mechanisms in the dorsal horn include enhanced synaptic strength and removal of inhibitory modulation, as well as microglial activation^11–13^. The critical outcome of this sensory plasticity is that an altered neuronal representation of tactile, thermal, or chemical stimuli is transmitted to the brain’s somatosensory circuits, contributing to heightened pain perception^14^.

The principal pathway by which nociceptive information reaches the brain is the anterolateral tract (ALT), a projection originating from a sparse subset of neurons within the dorsal horn. Despite its critical role in transmitting nociceptive information to the brain, the functional organization of the ALT remains incompletely characterized, both in protective and pathological pain states^15^. Recently, identification of marker genes in ALT neuronal populations has revealed that ALT neurons are molecularly and anatomically heterogenous^16–20^. Interestingly, while some transcriptionally defined subsets exhibit a narrow and homogeneous sensory tuning^17,19^, others appear to encode multiple stimulus modalities^16,18,21,22^. The molecular, functional, and anatomical heterogeneity of ALT projection neurons raises the possibility that diverse somatosensory dysfunctions such as inflammatory pain or neuropathic pain and chronic itch may be mediated by distinct perturbations in different sets of spinal outputs.

Given its critical role in transmitting nociceptive signals to the brain, altered signaling by ALT projection neurons must be a key component of clinical pain states. Indeed, acute electrophysiological studies have identified aberrant patterns of ALT activity in models of pain hypersensitivity^23,24^. Measurements of ALT neuron function have, however, so far been limited only to very acute time scales ranging from minutes to hours^25–27^, precluding longitudinal measurement of neuronal activity in the same set of neurons over the extended duration of clinically relevant models such as nerve-injury induced neuropathic pain^28,29^. It is therefore not known how the tuning of individual ALT neurons is altered or preserved during the induction and maintenance of persistent pain states, or whether nociceptive and neuropathic pain signals are transmitted from the spinal cord to the brain by the same or different ensembles of neurons. *In vivo* and *ex vivo* cellular imaging techniques can now be applied to the superficial dorsal horn of the spinal cord, allowing direct visualization of population sensory responses in spinal projection neurons and interneurons^26,30–36^. We have adapted this approach to allow repeated imaging cellular function *in vivo* over prolonged time scales, and applied it to track how sensory coding in spinal output circuitry is transformed during the development and maintenance of persistent neuropathic pain.

## RESULTS

### Longitudinal calcium imaging of superficial dorsal horn neurons

While a subset of ALT projection neurons are located in the deep lamina of the dorsal horn, particularly lamina V, the majority reside within the superficial lamina I, enabling optical access from the spinal surface for *in vivo* calcium imaging. We generated mice carrying either *Gpr83*^CreERT2^ or *Tacr1*^CreERT2^ together with a Cre-dependent GCaMP6f allele (Ai95)^37^. (**Figures 1A-B**). In these mice, GCaMP6f is expressed in the two genetically defined populations (*Gpr83*^+^ and *Tacr1*^+^ subsets), which include local interneurons and ALT projection neurons innervating diverse brain targets such as the lateral parabrachial nucleus (PBN_L_), which is the largest ALT projection in rodents,as well as periaqueductal gray nucleus (PAG), and thalamus^16^ **(Figure 1A**). To record specifically from spinal projection neurons, we also used a cohort of Ai95 mice in which an adeno-associated virus with retrograde tropism^38^ carrying Cre recombinase was injected bilaterally into the PBN_L_, resulting in GCamP6f expression in spinoparabrachial (SPB) neurons (**Figures 1A-B)**. It is possible that other subsets of projection neurons, such as spinomesencephalic neurons innervating the midbrain, are retrogradely labeled from the PBN_L_ injections because their axons pass through the injection site; we therefore refer to these as ‘retPbN’ rather than SPB neurons.

**Figure 1.**
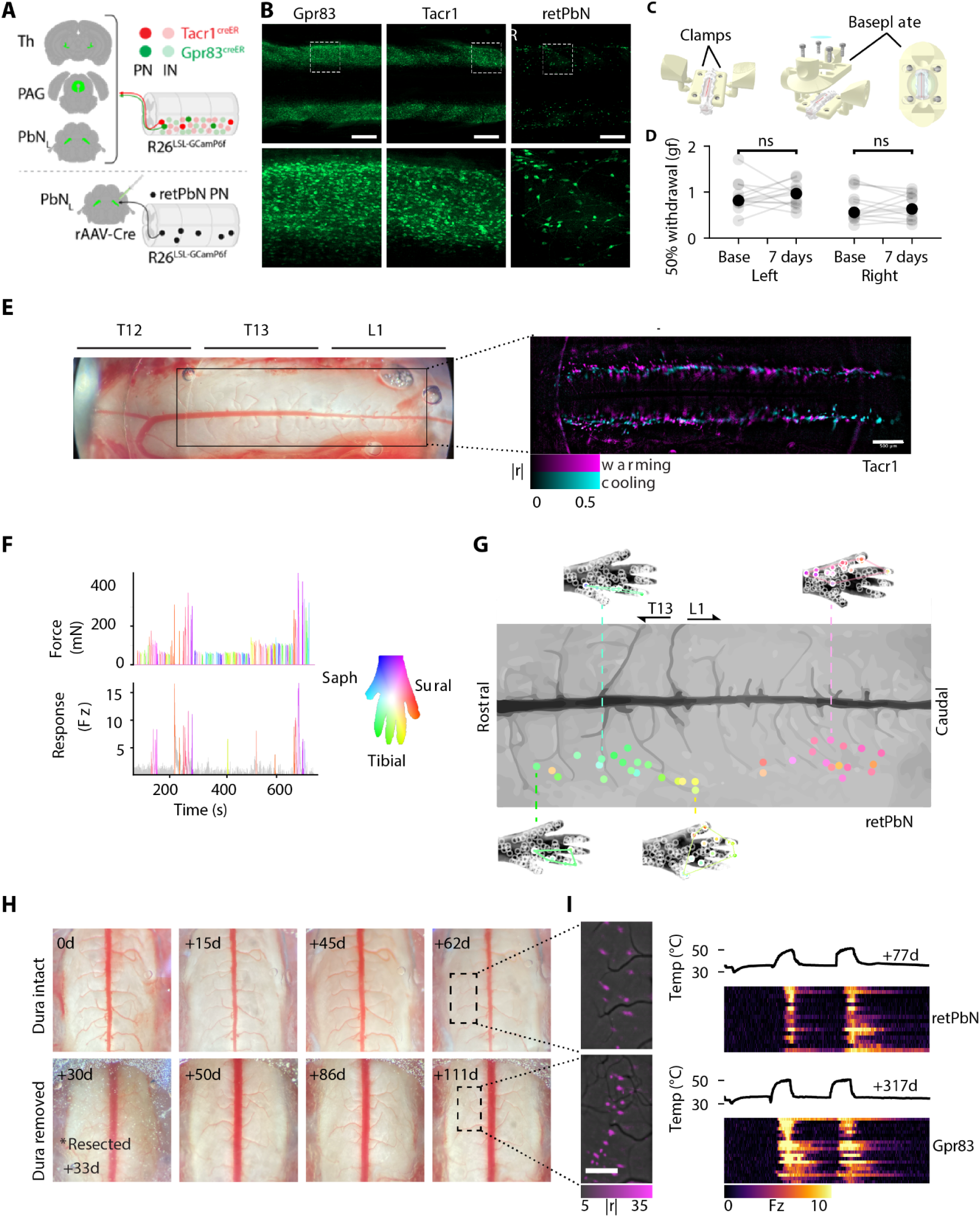
Longitudinal calcium imaging of superficial dorsal horn neurons. A. Targeting strategies to express the GCamP6f calcium reporter in dorsal horn lamina I projection neurons and interneurons. Top, animals carrying *Rosa^lsl-GCaMP6f^* allele and heterozygous for either *Tacr1*^CreERT2^ or *Gpr83*^CreERT2^ alleles label largely segregated sets of interneurons (INs) and projection neurons (PNs) projecting to posterior thalamus (Th), periaqueductal gray (PAG), lateral parabrachial nuclei (PBN_L_), and other ALT targets. Bottom, mice carrying the same Cre-dependent GCaMP6f allele are injected with a retrograde pseudotyped adeno-associated virus driving expression of Cre (rAAV-Cre), resulting in GCaMP6f expression exclusively in a subset of projection neurons. B. Maximal projections of cleared spinal cords expressing GCaMP, targeted anatomically or transgenically to superficial dorsal horn neurons. Maximum projections of GCaMP6f expression in superficial dorsal horn at low (top) and high (bottom) magnification are shown for transgenic *Gpr83^creERT^*^2^*; Rosa^lsl-GCaMP6f^* (left) and *Tacr1^creERT^*^2^*; Rosa^lsl-GCaMP6f^* (middle) animals, and for rAAV-Cre injected into *Rosa^lsl-GCaMP6f^* PbN (right). Scale bars 500 microns. C. Procedure for implantation of 3D printed window assembly. A laminectomy at the lumbar-thoracic vertebral level is clamped under a baseplate fixing a 5mm diameter coverslip over 1-3 spinal segments. The coverslip is sealed and adhered to the underlying tissue with a transparent silicone elastomer. D. Bilateral thresholds for 50% paw withdrawal thresholds to punctate mechanical stimulation (Von Frey assay) before and 1 week after the window implantation (p = 0.815 left, 0.680 right, paired t-test), indicating no postsurgical change in sensitivity. E. Micrograph of imaging window implanted above laminectomy at vertebral levels T12 to L1 of a *Tacr1^CreER^; Ai95* mouse, left, showing inset area containing neurons responsive to thermal stimulation of the paw. On right, composite of correlation maps shows pixel-wise absolute correlation of GCaMP signal to warming (magenta) or to cooling (cyan) stimuli across an imaging window. Correlation maps for each side of the spinal cord were generated from images acquired during stimulation of the paw ipsilateral to image acquisition. Scale bar 500 microns. F. Traces of force stimulation (Top), and z-scored GCaMP response intensity from an exemplary cell (Bottom), both color coded by location of indentation location on paw (right). Saphenous, tibial and sural nerves innervate the medial, central and lateral areas of the glabrous paw, respectively. G. Somatotopic map of receptive fields for mechanical indentation-responsive retPbN neurons. Paw maps illustrate the location of effective (filled) and ineffective (unfilled) stimuli. Each circle on the spinal cord image represents a single neuron, color-coded by the centroid of the receptive field according to the scheme shown in panel (E). H. Micrograph time series of spinal vasculature observed through windows at different time points after implantation. Top panels show a window with stably clarity after implantation. Bottom illustrates a window repaired and stabilized by resection of dura at 33 days post implantation. I. Stimulus-evoked GCaMP activity from the two windows shown in panel H, recorded 77 (Top) or 317 days (Bottom) after implantation. Maps on left show pixel-wise correlation to heating (bottom scale), rasters on right show temperature stimuli and rasters of GCaMP responses from imaged neurons. Color scale is Z-scored GCaMP fluorescence intensity (Fz).

As most behavioral assays and pain hypersensitivity models in rodents utilize the hindpaws^39,40^, we targeted our imaging to those lumbar spinal cord segments receiving input from the sciatic and saphenous nerves^41^. To achieve chronic optical access to these neurons *in vivo* we adapted elements of several previously reported designs^30,42^ to generate a 3D printed implant that stably fixes a 5mm diameter coverslip over a vertebral laminectomy at the lumbar-thoracic level (**Figure 1C**). The hindpaw behavioral mechanical sensitivity of spinal window implanted animals was normal one week after surgery as assessed by the withdrawal threshold to punctate stimulation of the plantar surface with Von Frey filaments (**Figure 1D**).

To evaluate sensory responses within the dorsal horn and characterize their somatotopic distribution we recorded GCaMP activity under moderate (1%) isoflurane anesthesia while delivering mechanical or thermal stimulation to the hindpaw (**Figures S1A,B**). Heating or cooling of the hindpaw evoked robust, spatially extensive, and stimulus selective fluorescence responses ipsilateral to the targeted paw **(Figure 1E, Supplemental Movie 1**), in a mediolaterally restricted zone extending from vertebral segments T13-L1, which correspond to spinal segments L3-L6. Punctate indentation of the glabrous paw skin elicited neuronal activity within the same area, with the rostrocaudal location of mechanically responsive neurons related roughly to the mediolateral location of their receptive field on the paw (**Figures 1F,G, Sup. Movie 2**).

To track the tuning properties of lamina I neurons in persistent pain states, imaging fields must be stable over periods of 3 weeks or longer to first establish baseline tuning and then identify changes in cellular responses over time. As noted in previous studies^42^, spinal windows frequently exhibit a degraded clarity within the first post-surgical week due to neovascularization and fibrosis over the imaging area, obscuring vascular landmarks and degrading or eliminating stimulus-evoked fluorescence signals from previously responsive regions **(Figure 1H**). However, by tracking the morphology of the spinal vasculature we identified a cohort of animals in which spinal windows stably maintained optical clarity for many months (**Figure 1**H). We further found that a subset of degraded windows could be rescued and stabilized for subsequent imaging by a single surgical resection of the dura over the imaging field. Using these stable and repaired chronic windows, robust stimulus evoked activity in a defined neuronal population could be imaged for months without further manipulation **(Figure 1I**).

### Cutaneous thermal information is encoded by three functionally distinct classes of ALT neurons

How are somatosensory stimuli represented in the population activity of lamina I dorsal horn neurons in healthy mice, and conveyed to post-synaptic targets in the brain by ascending projections? Previous electrophysiological and calcium imaging studies have demonstrated activation of projection neurons in superficial dorsal horn by thermal, mechanical and chemical stimuli^24,26,34,43^, yet it is unclear to what extent these responses are distributed broadly or selectively across different functionally defined cell types^44^. We therefore set out to define the population coding logic for thermal and mechanical stimuli in the healthy basal state.

Detection of innocuous temperatures allows animals to identify comfortable, safe thermal environments and maintain thermal homeostasis, while extreme temperature stimuli, particularly rapid heating, trigger defensive and escape behaviors. To probe responses to a wide range of thermal stimuli we stimulated the hindpaw of mice anesthetized with 1% isoflurane with a thermal spectrum consisting of rapid thermal transients from a neutral baseline of 34°C to temperatures ranging from 5 to 50 °C (see **Figure 1E**). Imaging of the ipsilateral dorsal horn revealed robust responses in ensembles of retPbN neurons that were elicited by intense heating or cooling, and smaller subsets that were responsive to moderate/innocuous warming or cooling stimuli (**Figure 2A**).

**Figure 2.**
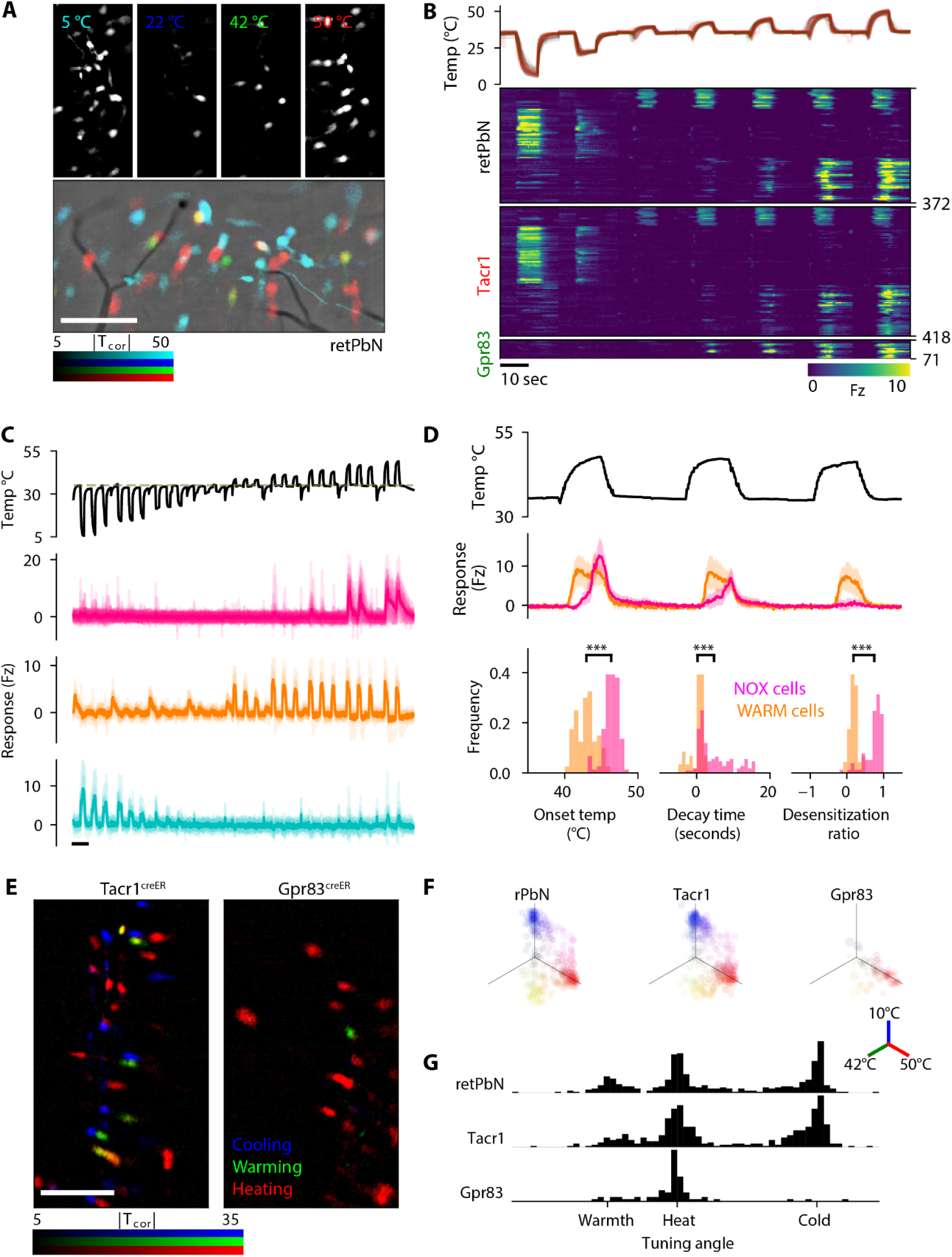
Three functionally distinct classes of ALT neurons encode cutaneous temperature. A. Maps of retPbN neuronal responses to intense (5 and 50 °C) and moderate (22 and 42 °C) cooling and heating. Images depict pixel-wise GCaMP fluorescence correlation with the thermal stimuli, individually (Top), and overlaid (Bottom). Scale bar 200 microns. B. Raster of GCaMP fluorescence responses to thermal spectra in naive mice, aligned to stimulus presentation and grouped by the three animal cohorts tested. Individual thermal stimuli are overlaid above. C. Example traces illustrating GCaMP fluorescence transients in superficial dorsal retPbN neurons for NOX (magenta), WARM (orange), and COLD (blue) categories of thermal responses. Solid traces are mean response for population, individual traces are overlaid in light color. D. Comparison of responses of WARM (orange) and NOX (magenta) neurons. Top panel shows temperature application, middle panel; mean traces of the two classes to repeated presentation of heating stimuli, ± SEM. Bottom panels show response onset, decay time relative to end of stimulus, and intensity of third response relative to first response (desensitization) for both classes (n=68 WARM, 133 HEAT cells, comparison by Mann-Whitney test). E. Overlaid correlation maps of responses to cooling(<10 °C), warming (42 °C) or heating (>50 °C) stimuli in Tacr1 and Gpr83 lamina I dorsal horn populations. F. Polar plots of thermal stimulus selectivity for retPbN, Tacr1 and Gpr83 neurons. Each marker represents the median normalized response of a single neuron to cooling, heating, and warming stimuli, summed between the three orthogonal axes (10, 42, and 50 °C). N = 372 retPbN cells, 418 Tacr1 cells, 71 Gpr83 cells. G. Histograms of thermosensory tuning angles calculated for cells in panel F. Angles labeled ‘Hot’ and ‘Cold’ correspond to exclusive responses to 50 °C or to 10 °C deg stimuli, ‘Warm’ corresponds to equal response amplitude to 42 °C and 50 °C stimulation.

A comparison of the ensemble responses to these different stimuli revealed that ALT thermosensory neurons are heterogeneous in their tuning. To identify and categorize the diversity of spinal thermosensory outputs we first aligned the responses of retPbN-GCaMP neurons to the thermal spectrum to generate a normalized response matrix, and then, in parallel, performed principal component analysis and K-means clustering to identify prototypical response classes **(Figures 2B,C**, **S2A**). Both analyses independently identified three major response types. The most abundant class (COLD cells) responded preferentially to cooling, with a maximal response elicited by stimuli below 10 °C. The remaining clusters were activated by primarily by increase in temperature, one that was maximally responsive at noxious temperatures (>50 °C, NOX cells) and one maximally responsive at or below the psychophysical threshold for heat induced pain (∼42 °C, WARM cells) in rodents and humans.

A comparison of the responses of WARM and NOX neurons to intense heating stimuli revealed differences in sensitivity, kinetics, and adaptation (**Figures 2D**). WARM neurons responded to heating transients with a shorter latency than noxious heat neurons and returned to baseline upon or prior to cessation of the heating stimulus. In contrast, NOX neurons commonly exhibited an after-discharge extending beyond removal of the heating stimuli. As a result, the population response to noxious heat was biphasic, with a sequential activation first of WARM neurons followed by the activation of the NOX neurons. The two populations were additionally distinguished by different desensitization properties; upon repeated heating to 47°C at 16 second intervals, WARM cells maintain consistent responses while nearly all NOX neurons were nonresponsive to a third stimulus. Together, these data indicate a functional segregation of innocuous and noxious heat responses in the mouse dorsal horn and suggest that thermal information transmitted through the ALT is organized into at least three distinct lines capturing cooling, warming, and noxious heat stimuli.The selectivity of neurons tuned to cold, warm and noxious heat stimuli was maintained both in recordings under 1% isoflurane anesthesia and in awake, behaving mice (**Figures S2C,D**)

We next examined the relationship between thermal tuning and the transcriptional identity of superficial dorsal horn neurons by recording responses to cold, warm and noxious hot stimuli in both Gpr83-GCaMP and Tacr1-GCaMP mice (**Figure 2E**). Despite the heterogeneity of these populations, which include spinoparabrachial and non-spinoparabrachial projection neurons, as well as spinal interneurons, we found that the thermosensory responses of these neurons encompassed the same three classes identified in retPbN neurons (**Figures 2F,G**). All three WARM, COLD and NOX type responses were represented in the Tacr1+ neurons, in similar proportions as in the retPbN population (**Figure S2B**). In contrast, Gpr83 neurons were preferentially selective for noxious heat (72% of neurons) or warmth, with minimal cold responses. Tacr1 expression defines, therefore, a population of interneuron and projection neurons in superficial dorsal horn that encompasses the full repertoire of retPbN thermal responses, while thermosensitive Gpr83 neurons are largely tuned to noxious temperature.

### NOX spinal projection neurons relay both thermal and mechanical modalities

We next asked how mechanosensory stimuli are represented within the Gpr83, Tacr1 and retPbN populations. We first characterized the mechanosensory input to the lumbar dorsal horn by imaging responses to mechanical stimulation of the plantar surface in Vglut2^cre^; Ai95 primary sensory neurons in the L4 dorsal root ganglia (DRG). Low-intensity (10-30mN) indentation and brushing reliably elicited calcium transients in numerous L4 DRG neurons (**Figures 3A,S3A**) In contrast, responses to the same tactile stimuli were only observed infrequently in the retPbN neurons of the superficial dorsal horn (**Figures 3A,S3B**). Both DRG and retPbN neurons responded robustly to high-intensity (>100mN) indentation of the plantar surface (**Figure 3A**),

**Figure 3.**
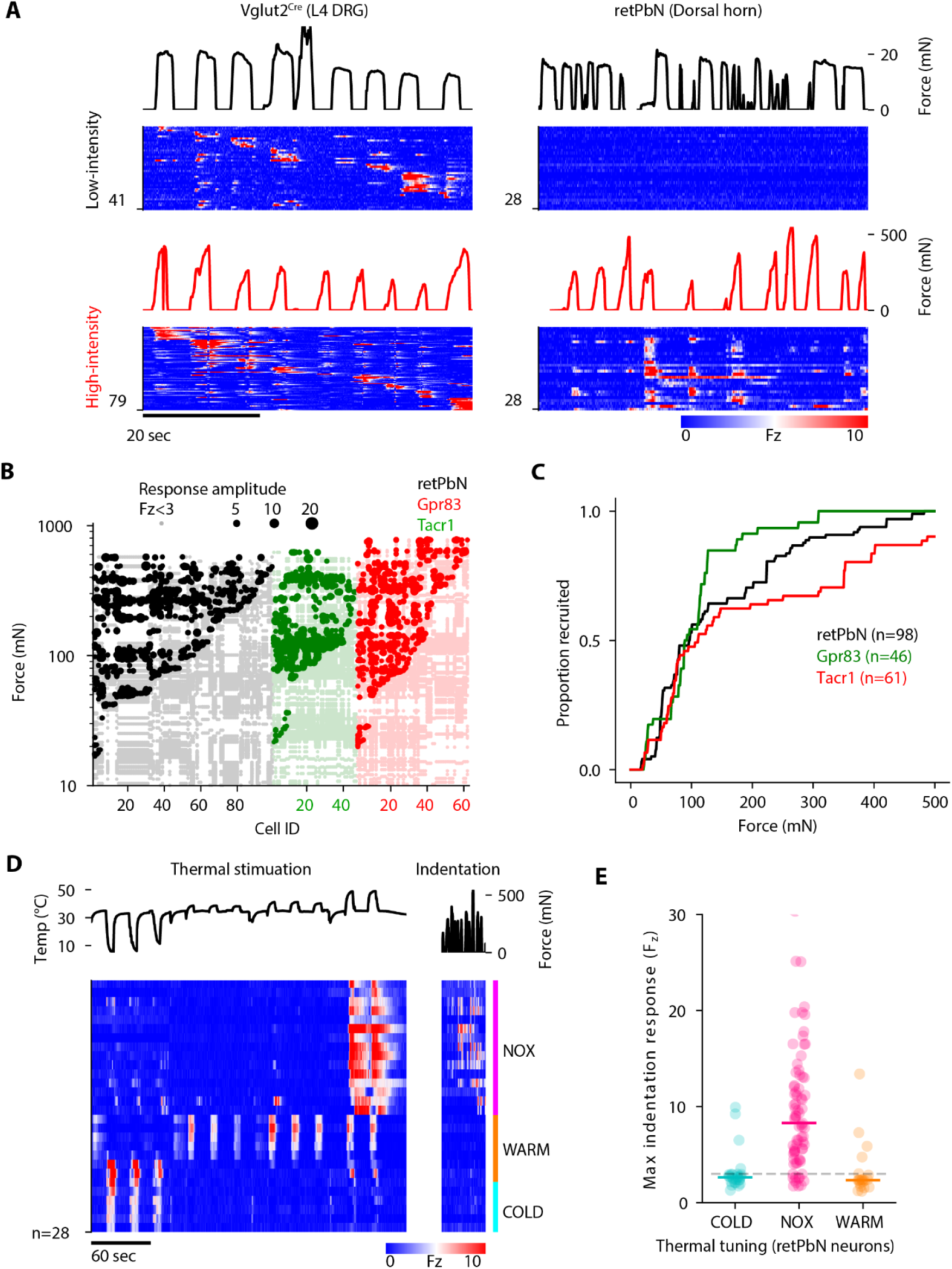
Nociceptive ALT neurons encode high-intensity mechanical stimuli. A. Rasters of mechanically evoked GCaMP signals recorded from *Vglut2^cre^* lumbar DRG neurons (left), and retPbN dorsal horn neurons (right), in response to low-intensity indentation (10-25mN, top), or moderate to high-intensity indentation (100-500mN, bottom). B. Plot of GCaMP responses evoked in retPbN, Tacr1 and Gpr83 neurons by indentation. Individual cells are plotted on x axis, each circle corresponds to a single stimulation, marker size corresponds to response amplitude above threshold (Fluorescence z-score ≥ 3 for 200ms). Stimuli not evoking responses are plotted with light color marker. C. Cumulative recruitment of mechanically sensitive neurons for each of the three lamina I dorsal horn populations with increasing indentation force. N = 10 mice. D. GCaMP fluorescence raster of retPbn neurons stimulated with thermal and mechanical stimuli, sorted by the three thermosensory response classes. E. Maximal response (z-scored fluorescence, Fz) of retPbN spinal projection neurons evoked by mechanical indentation of hindpaw skin, plotted neurons in each of the three functionally defined thermosensory classes. (NOX N = 68, COLD N=35, WARM = 21 cells from 4 animals).

To quantify the distribution of mechanical sensitivities of retPbN, Gpr83 and Tacr1 neuronal populations we presented a graded series of indentation stimuli across the plantar skin (**Figure 3B,C**). The median threshold to plantar stimulation did not vary significantly between these groups, ranging from 90.5 to 110.3mN (**Figure S3C**). Notably, retPbN neurons were significantly more sensitive to indentation of the hairy dorsal surface of the paw, with a median threshold of 54.9mN (**Figure S3C**).Behaving mice stimulated on the plantar surface with the same indentation probes responded with 50% withdrawal frequency to forces in the 30-40mN range(**see Figures S1C,D**). Thus, the majority of retPbN neurons were responsive only to mechanical stimulus intensities that reliably trigger avoidance behaviors in awake mice.

Next, we examined the extent and nature of the relationship between mechanosensory and thermosensory responses in ALT neurons. We first classified the tuning of retPbN neurons to thermal stimulation of the hind paw, and then probed the surface of the paw for mechanical responses (**Figure 3D**). Notably, we did not identify any cells solely tuned to mechanical stimuli, as all mechanically responsive neurons, including those responsive to innocuous stimuli (see **Figure S3A**), responded to at least one type of thermal stimulus. Mechanical sensitivity was differentially distributed across the three thermosensory classes, with the majority of mechanically responsive neurons (65/74, 88.7%) being members of the NOX thermosensory response cluster (**Figure 3E**). The NOX neuron population identified by responsivity to intense heat exhibited widespread mechanosensitivity, with 84.4% (65/77) responding to mechanical stimulation of the paw. These data support a coding scheme for ALT spinal output in which distinct cold and warm tuned populations narrowly convey innocuous thermal information, while a separate set of broadly tuned neurons signal the presence of both thermal and mechanical noxious stimuli.

### Peripheral sensitization alters the tuning of nociceptive ALT neurons

We next asked if and how the somatosensory responses observed in the health/naive state where pain is only evoked by noxious stimuli (acute nociceptive pain) are modified in conditions that produce acute pain hypersensitivity. To do this we examined the effect of sensitizing primary nociceptors by a brief application of a low dose (0.1%) topical capsaicin to the surface of the hindpaw (**Figure S4A**). The treatment elicited a robust behavioral hypersensitivity to heating that appeared within minutes and was fully resolved by 24 hours (**Figure 4A**). This approach enables analysis of the effect of direct sensitization of primary nociceptors without concomitant tissue damage such as that caused by an injection, burn, or incision^40^. Imaging of primary sensory neurons in the L4 DRG revealed that responses to warming were dramatically increased after the capsaicin treatment, confirming a robust induction of peripheral sensitization in this model (**Figure S4B**).

**Figure 4.**
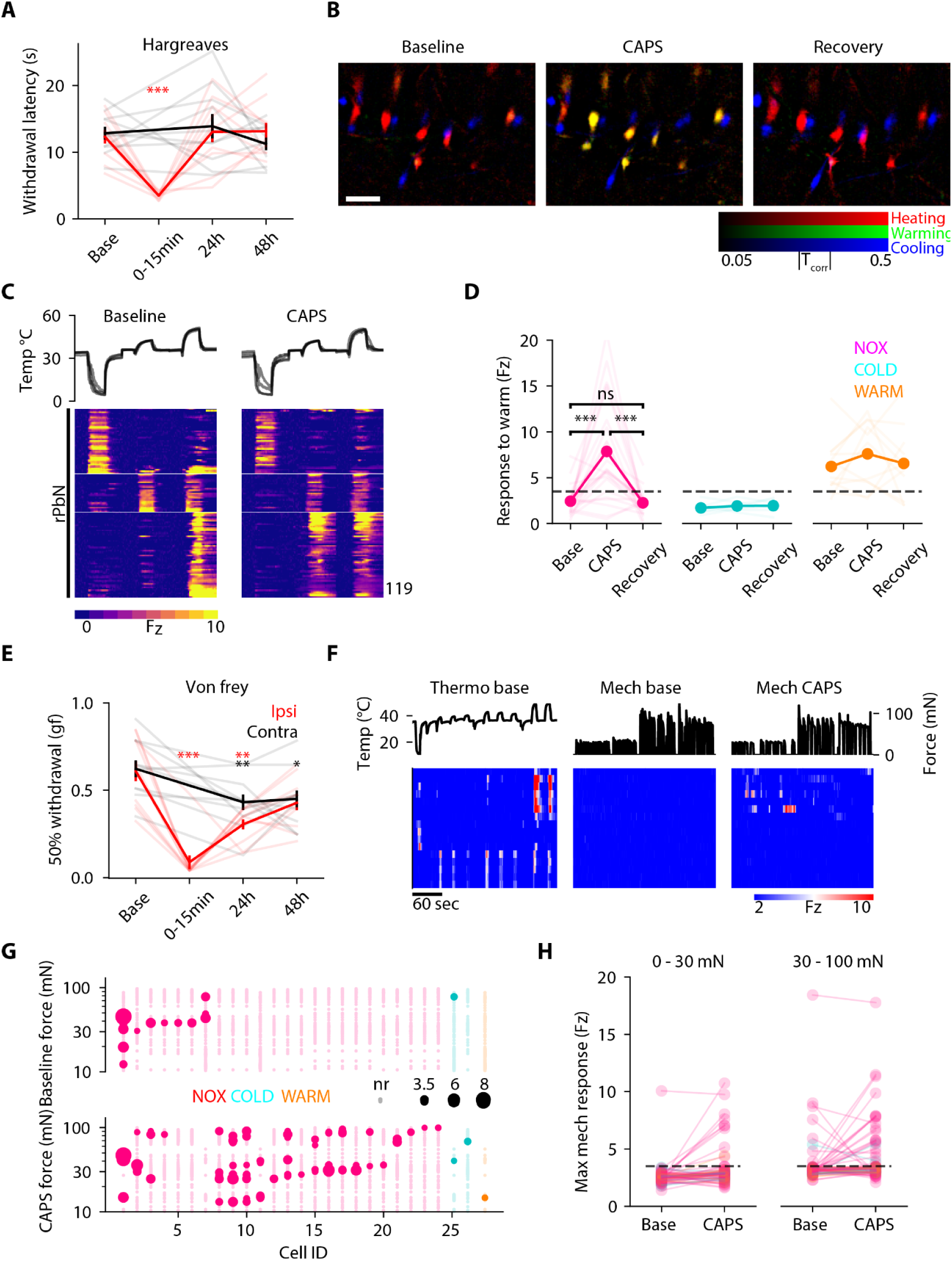
Topical capsaicin alters tuning of ALT nociceptive neurons to thermal and mechanical stimuli. A. Latency of ipsilateral and contralateral paw withdrawal to radiant heating (Hargreave’s test) in mice before and after topical low concentration capsaicin application to one paw. Immediate (0-15 minute) time point was recorded only for ipsilateral paw. (Dunnett’s multiple comparison to baseline value, N=10 mice. B. Overlaid maps of GCaMP signal correlation to cooling (<10 °C), warming (42 °C) or heating (>50 °C) recorded from an aligned retPbN imaging field prior to (Baseline), immediately after (CAPS), and 2 weeks after (Recovery) application of topical capsaicin to the ipsilateral hindpaw. Scale bar = 100 microns. C. Rasters of retPbN GCaMP responses aligned to cooling, warming, and heating thermal stimuli before (Baseline, left) and directly after (CAPS, right), topical capsaicin treatment. Recordings are from N=4 retPbN animals. D. Maximal responses of NOX, COLD, and WARM thermosensory neurons to a 42° C warming stimulus before, immediately after, and 2-3 weeks following topical capsaicin treatment (Dunn’s multiple comparison’s test, N = 26 NOX, 8 COLD, 13 WARM cells). E. Mechanical stimulus intensity eliciting 50% paw withdrawal (grams force, Von Frey test), evaluated before and after ipsilateral administration of topical capsaicin. Dunnett’s multiple comparison, N=11 mice. F. Rasters of GCaMP responses recorded from retPbN neurons in response to thermal stimulation and low to moderate intensity mechanical indentation before (Base) immediately following topical capsaicin treatment (CAPS). Stimulus intensity was limited to <200mN to avoid mechanical damage to skin. G. Plots of mechanical indentation-evoked calcium responses in retPbN cells color-coded by thermosensory class, before (Top) and after (Bottom) topical capsaicin treatment. Marker size correlates to response amplitude, stimuli not evoking responses (Fz<3) shown in light color. Recordings from 27 mechanically responsive cells in 3 ret PbN animals. H. Responses to low-intensity mechanical indentation appear in NOX neurons immediately after capsaicin treatment. Maximum z-scored GCaMP fluorescence amplitudes evoked by low (0-30mN, left) and moderate intensity (30-100mN, right) mechanical stimulation for retPbN cells before and after (15-30 minutes) topical capsaicin, color coded by basal thermosensory tuning of cells. N = 50 NOX cells, 14 COLD cells and 12 WARM cells..

In the dorsal horn, we observed a large expansion in the ensemble of ALT neurons responsive to innocuous warmth after the application of the topical capsaicin (**Fig 4B**). This increase in warmth responses manifested in a selective sensitization of the NOX population, which is normally tuned to noxious heat (**Figures 4C,D,S4C**). In contrast, the responses of cold and warmth sensing populations were not altered by the capsaicin treatment (**Figures 4C,D**). Consistent with the rapid recovery of the paw-withdrawal behavioral responses, the patterns of increased activity of the sensitized neurons returned back to baseline levels when recorded at later time points (**Figures 4B,D**). The sensitivity of the NOX ALT population to innocuous warming stimuli matches, therefore, both the appearance and the resolution of nocifensive reactions to warming in mice subjected to acute transient peripheral sensitization.

In addition to thermal allodynia, the topical capsaicin also rapidly elicited a robust hypersensitivity to punctate tactile stimulation of the hindpaw (**Figures 4E**). To examine the effect of topical cutaneous capsaicin on mechanosensory output from the dorsal horn, we first determined the thermal tuning of retPbN ALT neurons, and also recorded the activity elicited by low and moderate intensity indentation of the plantar skin. We then applied capsaicin and recorded responses to a second series of indentation stimuli (**Figure 4F**). A subset of neurons displayed enhanced responses to mechanical indentation after the capsaicin, including to low intensity (<30mN) stimuli that rarely evoke activity in retPbN neurons at baseline **(Figures 4F-H**). Responses to mechanical stimulation after capsaicin remained almost entirely restricted (21/24, 87.5%) to those superficial dorsal horn neurons belonging to the NOX thermo subset (**Figures 4F,G**). These data suggest that the acute behavioral hyper-reactivity to punctate mechanical stimuli observed after topical capsaicin may be attributed to a re-tuning of polymodal high-threshold (NOX) ALT neurons to low-intensity mechanical stimulation.

### Lamina I sensory tuning is stable over weeks in the absence of injury

Clinically relevant pain conditions typically affect patients over weeks or longer. To evaluate potential changes in spinal sensory tuning associated with persistent pain states, we first characterized the normal dynamics of stimulus-evoked activity in the dorsal horn on this time scale. Although innate responses to somatosensory stimuli are stable in the absence of injury^45^, it is not known whether the sensory tuning of individual dorsal horn sensory neurons is stable or changes over time. We therefore conducted longitudinal imaging experiments to characterize the long-term dynamics of spinal sensory tuning in healthy animals.

We imaged neuronal responses to sets of thermal stimuli over intervals of 1 to 6 weeks for retPbN, Gpr83 and Tacr1 populations. Stimulus evoked GCaMP response maps remained highly stable over these intervals (**Figure S5A**). Furthermore, the thermal tuning of individual neurons remained strikingly consistent over periods of days and weeks (**Figures S5B,C**). These data indicate that the stability of innate responses to somatosensory stimuli in healthy animals corresponds to stability at the level of cellular stimulus selectivity within superficial dorsal horn, including ascending thermosensory and nociceptive ALT projection neurons.

### Peripheral nerve injury alters innocuous thermosensory output from the dorsal horn

We next examined the extent to which the stable sensory tuning of the superficial dorsal horn is modified in a persistent, maladaptive pain state. Spared nerve injury (SNI) is a model of nerve injury-induced neuropathic pain that reliably induces robust pain hypersensitivity to thermal and mechanical stimuli, which persists through the lifetime of the animal. We first characterized stimulus-evoked GCaMP activity in neuronal populations of naive Tacr1, Gpr83, and retPbN mice with chronically implanted spinal windows, next we induced SNI by transecting and ligating the tibial and peroneal branches of the sciatic nerve (**Fig S5D**), and then imaged the same sets of neurons 2-3 weeks after the injury, when behavioral hypersensitivity to stimulation of those parts of the hindpaw innervated by the spared sural and uninjured saphenous nerves plateaus^28,46^.

As expected, all SNI mice with implanted spinal windows exhibited mechanical allodynia of the sural receptive field ipsilateral but not contralateral to the nerve injury when tested 4 weeks post-surgery (**Figure S5E**).A comparison of thermally evoked population responses before and after SNI in lamina I Tacr1 neurons revealed widespread changes in sensory tuning in the ipsilateral dorsal horn (**Figure 5A)**. In contrast, thermosensory tuning remained stable in neurons contralateral to the injured nerve (**Figures S5F-H**).

**Figure 5.**
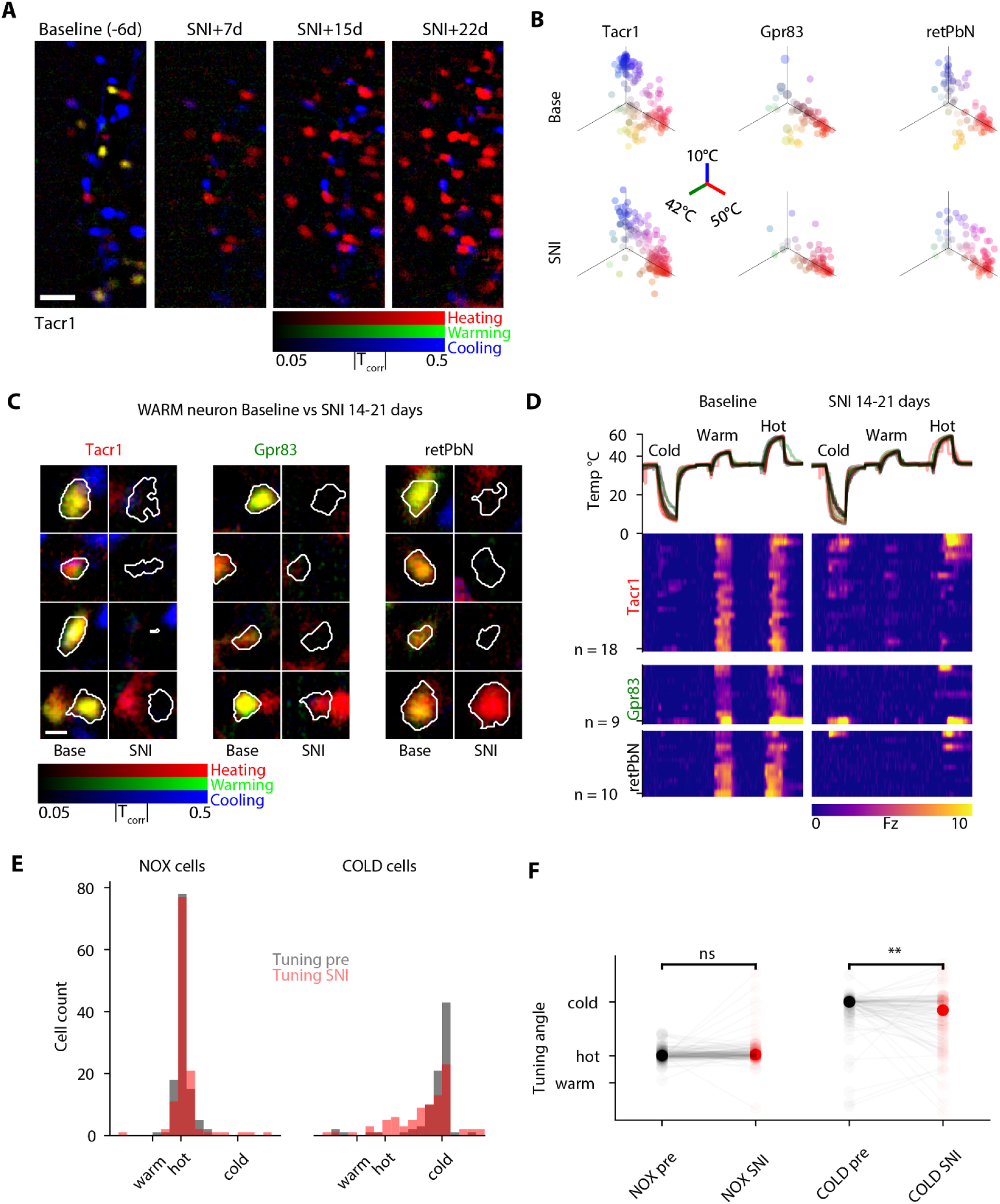
SNI modifies spinal representation and transmission of innocuous temperature responses. A. Responses to cooling (<10 °C, blue), warming (42 °C, green) or heating (>50 °C, red) recorded from an aligned Tacr1 imaging field are overlaid before (Baseline) and 7, 15, and 22 days after spared sural nerve injury. Scale bar 100 microns. B. Polar plots of thermal stimulus selectivity for sets of retPbN, Tacr1 and Gpr83 neurons at baseline (Top) and 2-3 weeks after SNI (Bottom). Each marker represents the median normalized response of one neuron to the cooling, heating, and warming stimuli, summed along the orthogonal axes. N = 81 retPbN cells, 161 Tacr1 cells, and 65 Gpr83 cells. C. Maps of GCaMP correlation to cold (<10° C), warming (42° C) and hot (50° C) temperature for neurons tuned to WARM in naive mice (Base), and correlation maps of the same imaging location recorded 2-3 weeks after ipsilateral SNI, for Tacr1, Gpr83, and retPbN neurons. Contours of the regions used to extract calcium signals are overlaid for each image. Scale bar 25 microns. D. Rasters of GCaMP responses of WARM responsive Tacr1, Gpr83, and retPbN neurons aligned to cooling, warming, and heating thermal stimuli before (Baseline) and 14-21 days after SNI. GCaMP activity recorded from N=4 Tacr1, 3 GPr83 and 4 retPbN mice. E. Overlaid histograms illustrating distribution of thermosensory tuning angles for the NOX and COLD subpopulations measured at baseline before SNI (grey) and in the same neurons measured 2-3 weeks after SNI (red). F. Paired plots of thermosensory tuning angles for each COLD and NOX class neuron before and after SNI. Compared by paired t-test, N = 87 COLD cells, 120 NOX cells)

An examination of the distribution of thermal tuning among Tacr1 and Gpr83 neurons, which together constitute the majority of spinoparabrachial projection neurons, as well as superficial interneurons, revealed that WARM neuronal responses were absent from these populations after SNI (**Figure 5B**). Furthermore, WARM responses were also eliminated after SNI in retPbN spinal projection neurons (**Figure 5B)**. Tracking of individual WARM neuron responses demonstrated that although a subset retained sensitivity to noxious heating stimuli (>50° C), the majority did not respond to any thermal stimulus after SNI (**Figures 5C,D**). These data indicate that after SNI there is essentially a complete loss of transmission between the superficial dorsal horn and PbN_L_ of innocuous warming signals from the affected hindpaw.

In addition to profound loss of WARM cell function, we noted an increase in the population of neurons responsive to both noxious heat and cold within the retPbN and Tacr1 populations, but not in Gpr83 neurons (**Figure 5B**). To investigate the source of this new polymodal response, we characterized the effect of SNI on neurons in both the basal COLD and NOX populations. While basal NOX neurons maintained a narrow tuning to noxious heat stimuli after SNI (**Figures 5D,E**), the COLD population exhibited a decreased selectivity for cooling, with a subset now responding to both cooling and noxious heating stimuli **(Figures 5D,E**).

The tuning of NOX neurons present in the naive, pre-injury state was largely unchanged after SNI (**Figures S6A,B)**. In contrast to the sensitization of NOX neurons to innocuous thermal and mechanical stimuli induced by topical capsaicin treatment, we did not observe any increased sensitivity to low intensity warming (**Figures S6A,B**). Similarly, while mechanical allodynia was induced in all mice ipsilateral to SNI **(Figure S5E**), responses to low-intensity mechanical stimulation (<30mN) remained rare across the retPbN, Gpr83 and Tacr1 neuronal populations (**Figures S6C,D**). The median mechanical response thresholds of Gpr83 and retPbN populations did not change significantly. Tacr1 neurons exhibited moderately lower thresholds to mechanical indentation, but did not respond to low-intensity stimuli. High-intensity indentation remained an effective stimulus for the majority of NOX neurons **(Figures S6C,D**).

### Silent neurons are recruited persistently in chronic in neuropathic pain

Despite the stability of mechanical and noxious heat responses in NOX neurons after SNI, we observed a striking expansion in that set of neurons activated only by noxious temperatures. An alteration of tuning in basal COLD and WARM neurons into a NOX type response was insufficient to account for this increase (see **Figures 5D,F**). We discovered cells in both the Tacr1 and Gpr83 populations that did not respond to any stimuli in the naive state, but which acquired sensitivity to intense heat after SNI, which we classified as ‘SILENT’ neurons (**Figures 6A,B**). As Tacr1 and Gpr83 populations include both interneurons and projection neurons, we next asked if the SILENT neurons contribute to ALT output after SNI, and found a set of SILENT neurons within the anatomically defined retPbN projection neuron population (**Figures 6A-B**).

**Figure 6.**
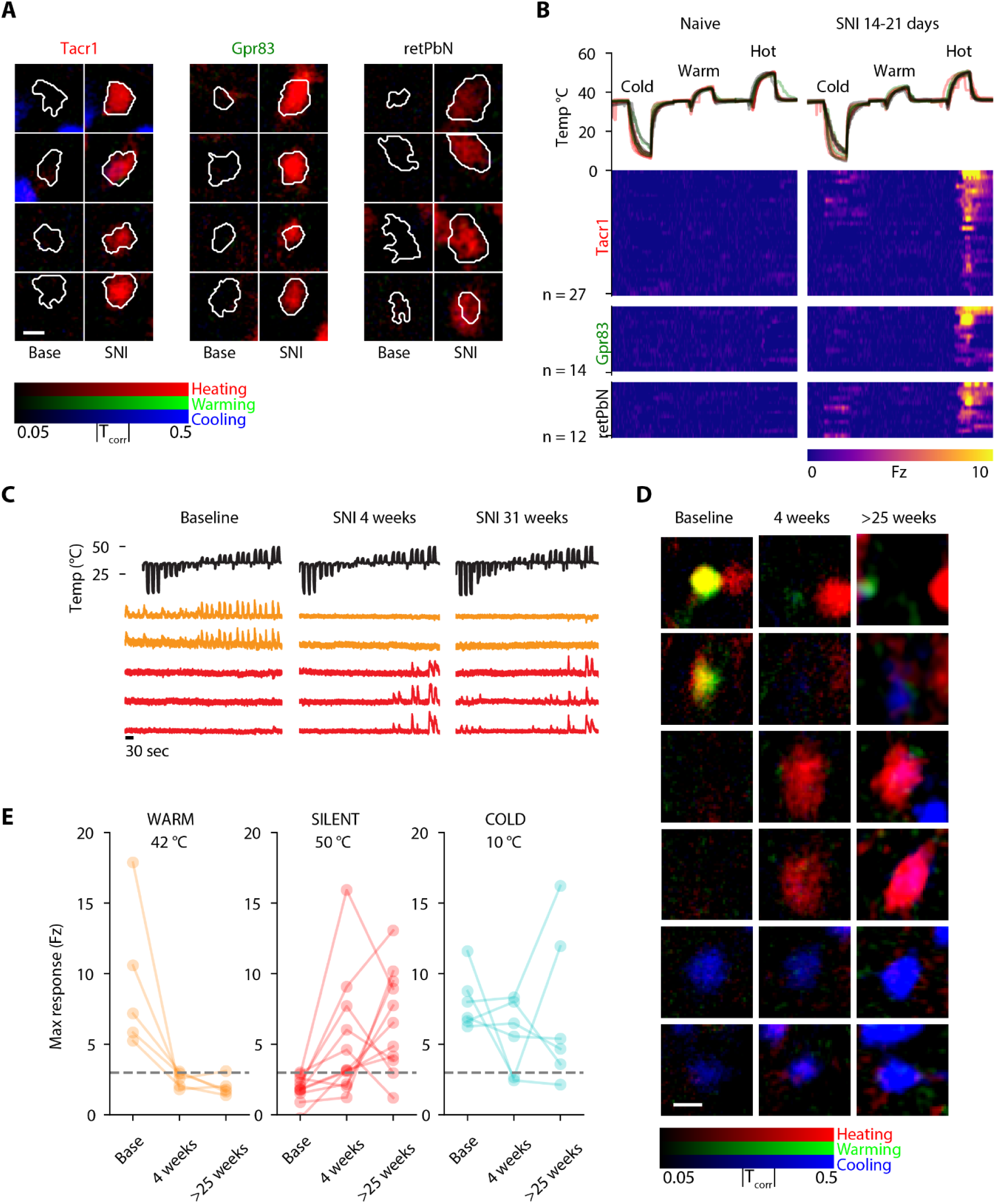
Recruitment of SILENT neurons in neuropathic pain. A. Correlation maps between GCaMP fluorescence and thermal stimuli for SILENT neurons in Tacr1, Gpr83, and retPbN populations. Paired images show imaging locations aligned between sessions recorded before and 14-21 days after ipsilateral SNI, for Tacr1, Gpr83, and retPbN populations. Contours of regions used to extract calcium signals are overlaid for each image. Scale bar 25 microns. B. Rasters of GCaMP fluorescence intensity for SILENT class Tacr1, Gpr83, and retPbN neurons aligned to cooling, warming, and heating thermal stimuli before (Base) and 14-21 days after SNI. GCaMP activity recorded from N=4 Tacr1, 3 GPr83 and 4 retPbN mice. C. GCAMP fluorescence traces for WARM (orange) and SILENT (red) neurons recorded before SNI (Baseline), at 4 weeks after SNI, and at >25 weeks after SNI. D. Correlation maps for exemplary WARM and SILENT neurons tp thermal stimuli imaged at baseline, 4 weeks, at >25 weeks after SNI.Scale bar is 25 microns. E. Maximum z-scored GCaMP fluorescence response amplitudes elicited by preferred stimulus at baseline, 4 weeks, and >25 weeks after SNI for WARM, SILENT, and COLD neurons. GCaMP activity recorded from N=2 Gpr83, and 2 retPbN mice.

The neuropathic pain state induced by SNI persists well beyond the 2-3 week period in which we identified an activation of SILENT neurons and loss of innocuous thermosensitivity. The chronic stage SNI (>3 months) can be distinguished from the acute phase by long-term plasticity of both peripheral and central neurons^12,29^. In a subset of Gpr83 and retPbN mice we imaged responses to temperature >25 weeks after SNI, well into the chronic neuropathic pain phase. Remarkably, we found that both the loss of responses in WARM cells, and the activation of SILENT neurons, persisted at this time (**Figures 6C-E**). These data suggest that alterations in the output from the dorsal horn established in the acute phase of nerve-injury induced neuropathic pain remain present in the chronic phase.

### Plasticity of distinct spinal outputs underlies acute sensitization and persistent neuropathic pain states

Acuteperipherally sensitized and neuropathic pain states, are associated then, with clear and distinct modifications in the sensory tuning of superficial spinal projection neurons. To visualize and compare changes in thermosensory function across these distinct pain states we constructed fate maps tracking the tuning of individual neurons over time. These indicate that the population representation of thermosensory stimuli transmitted from the spinal cord to the brain via the ALT is stable in healthy conditions (nociceptive pain elicited only by noxious stimuli) (**Figure 7A**), but undergo a significant and stereotyped reorganization upon the transition to states where pain is elicited by normally innocuous stimuli **(Figures 7B,C**).

**Figure 7.**
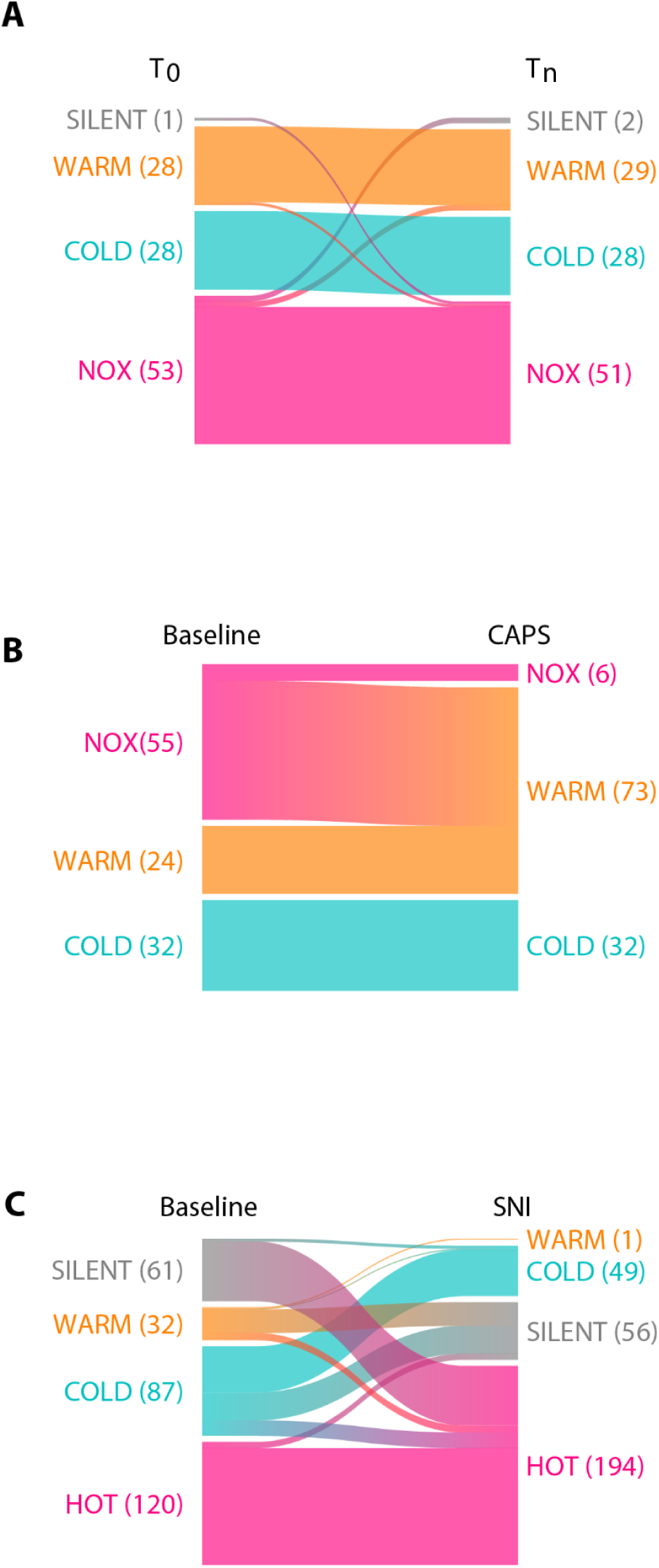
Transformations of sensory tuning in acute and persistent pain. A. Sankey plot depicting transitions of neurons between functionally defined thermosensory profiles over two imaging sessions, in the absence of any treatment. Responsive cells are classified according to thermal tuning preference, neurons are classified as ‘none’ if no temperature evoked calcium transients were observed within spatial footprint. Diagram collates data from retPbN, Tacr1, and Gpr83 populations. B. Alteration of basal thermosensory tuning (left) after topical capsaicin treatment (right). All recordings were made immediately before and immediately after (1-30 minutes) treatment. All cells are spinal projection neurons (retPbN). Note transformation of ‘heat’ tuned neurons to warmth sensitive class. C. Transitions in therrmosensory tuning observed 2-3 weeks after SNI. Note appearance of heat selectivity in cells unresponsive cells (SILENT) at baseline and loss of response in WARM neurons. Diagram collates data from retPbN, Tacr1, and Gpr83 populations.

The plasticity in ALT tuning induced by the peripheral sensitization of TRPV1 expressing nociceptors by topical capsaicin is primarily a specific modification of function within the NOX population, as COLD and WARM neurons largely retain their basal tuning (**Figure 7B**). The majority of NOX neurons acquire sensitivity to normally innocuous warming stimuli after topical capsaicin, which aligns with the aversive behavioral responses to such stimuli evoked after this treatment and the reduction of threshold of heat sensitive nociceptors at the periphery. Together with the sensitization of NOX neurons to low-intensity mechanical stimulation, these data suggest that the acute pain hypersensitivity induced by a peripherally-acting sensitizer is accompanied by a re-tuning of normally high-threshold polymodal ALT neurons to low-intensity thermal and mechanical stimuli, allowing these innocuous stimuli to engage nocifensive responses and pain perception in the brain.

In contrast, SNI results in a quite different transformation of thermosensory representations in the superficial dorsal in multiple functionally defined cell types. The conservation of NOX responses, along with the *de novo* recruitment of heat responses in a normally SILENT population and the reduced selectivity of COLD neurons results in an expansion in the ensemble of superficial dorsal horn neurons responsive to noxious heating (**Figure 7C**). Together with a loss of WARM cell function, these changes shift the balance of activity in ALT and superficial dorsal horn neurons towards noxious heat nociceptive signaling in a sustained manner for many months.

## DISCUSSION

### Normal coding of thermal and mechanical stimuli in the ALT

In this study longitudinal *in vivo* calcium imaging of specific sets of superficial lumbar dorsal horn neurons was applied to both characterize the functional organization of the ALT and track its modification over time in different pain models. In the basal nociceptive state where only an acute noxious stimulus evokes pain, we reveal the general logic of the normal encoding of somatosensory stimuli, one that incorporates both labeled-line and integrative coding. We find that high-intensity thermal and mechanical stimuli converge on a common population of superficial dorsal horn neurons (NOX) that have, in consequence, a broadly-tuned nociceptive output. In contrast, cooling and innocuous warmth are transmitted to brain somatosensory circuits by segregated, narrowly tuned populations that display stable tuning in the absence of major perturbations. These three different populations reflect the ethological contribution of discriminative temperature sensing of temperature gradients to both identify a thermal comfort zone and initiate escape and avoidance behaviors in response to imminent danger/noxious stimuli regardless of the stimulus modality^47,48^.

While cold selective superficial dorsal horn neurons, including spinoparabrachial projection neurons, have been previously characterized^49^, warm sensitive neurons were encountered only infrequently in previous studies of spinal sensory tuning^50^. However, consistent with our identification of a population of warm sensitive spinal projection ALT neurons, functionally and transcriptionally defined populations of cells with similar thermal responsiveness are present in the mouse PbN^51,52^. Together with warm-sensing neurons in the insula^53^, these findings support the existence of a dedicated pathway for warmth detection from the spinal dorsal horn to the brainstem and cortex. An analysis of afferent and interneuronal inputs to the ALT warm neurons would aid identification of which afferents are responsible for the primary sensing of innocuous warmth, which, so far, remains elusive^54–56^.

Our analysis was restricted to thermal and mechanical somatosensory inputs arising from the hindpaw, it is likely that exploration of additional stimuli and body areas may identify lamina I ALT populations that convey pruriceptive, viscerosensory, or affective touch signals to the brain^17,57,58^. We did identify small subsets of retPbN, Gpr83 and Tacr1 lamina I neurons responsive to low-intensity mechanical stimuli. While recordings from spinoparabrachial neurons in anesthetized mice and rats indicate that the majority are only sensitive to intense or noxious stimuli, analyses in skin-spinal cord preparations suggest that most spinoparabrachial neurons innervating hairy skin are responsive to light or moderate tactile stimuli when recorded *ex vivo* in the absence of anesthesia or descending inputs. Our recordings from awake behaving mice suggest that the temperature tuning of lamina I neurons is largely conserved under 1% isoflurane anesthesia, however we were not able to accurately quantify the mechanical forces acting upon hindlimbs in the awake state sufficiently to perform a similar comparison for mechanical stimuli. Recording of lamina I responses in awake mice^30,59^, combined with well-controlled quantitative mechanical stimulation, is needed to assess the physiological and behavioral relevance of spinal mechanosensory responses measured under varying conditions.

### Re-mapping of ascending thermosensory representations in acute and chronic pain

Upon the induction of a peripheral sensitized pain state, each functional class of lamina I sensory neurons exhibits stereotyped patterns of plasticity. How do these changes relate to the behavioral and perceptual dysfunctions observed in pathological pain? In principle, alterations in information transfer from the spinal cord to the brain may contribute to perceived pain hypersensitivity through increased the sensitivity or activity of existing nociceptive projections, recruitment of additional nociceptive populations, or by emergence of aberrant patterns of activity across multiple populations^14^.

Our data indicate that the emergence of responses to low-intensity stimuli by the broadly tuned nociceptive projection pathways may contribute to the perception of thermal hypersensitivity following acute peripheral nociceptor sensitization. This mechanism does not appear, though, to drive the allodynic sensory phenotype of neuropathic pain. Instead, we find that a loss of innocuous sensory outputs, and an activation of a previously silent pool of spinal projection neurons, constitute the circuit motifs that may contribute to the specific hyperalgesic elements of neuropathic pain. These results indicate that pain hypersensitivity mediated by acute nociceptor sensitization or by peripheral nerve injury are associated with different modifications of sensory tuning in entirely distinct populations of spinal ALT output neurons, and with very different durations (hours versus months).

The changes in the tuning of spinal output neurons are concordant with behavioral features of the pain models examined. The reversible hyperreactivity of NOX cells to warming stimuli after topical capsaicin aligns with the robust but transient heat allodynia exhibited by awake behaving animals receiving the same treatment. Likewise, animals with SNI exhibit heat hyperalgesia but not warmth allodynia^28^, consistent with an expansion of the neuronal response to noxious heat by recruitment of SILENT neurons and the loss of WARM neuron function.

The complete loss of sensibility to innocuous warmth in superficial dorsal horn projection neurons following SNI was an unanticipated feature of the neuropathic pain state. This selective loss could be explained either by a specific loss of sensitivity of warmth-sensitive primary afferent neurons in this injury model, or by a disrupted spinal processing of normal thermal inputs. If input from cold-selective primary afferents is a salient signal for warmth detection^56^, the preservation of COLD responses in ALT neurons after SNI would suggest that loss of warmth sensation is mediated by spinal mechanisms post-synaptic to primary sensory neurons. Alternatively, if warmth is detected by dedicated warmth sensors in the DRG^54^, warmth sensing neurons in the spared sural and saphenous nerves may alter their sensory function in response to damage of the tibial and peroneal nerves at the periphery by signaling from injured neurons or immune cells in the DRG.

What is the potential significance of an altered representation of innocuous stimuli to pain sensation? Somatosensory signals of distinct modalities are relayed in parallel from the spinal cord to brainstem and thalamic targets, where they undergo further processing before eliciting perceptual, affective, autonomic, and motor responses. For example, a spatial interleaving of innocuous warmth and cold stimuli can produce an illusion of pain in human subjects^60^, and deficits of innocuous somatosensation manifest in central pain syndromes^61^. The altered representation of somatosensory stimuli we observe in the SNI neuropathic pain model will result in patterns of activity across multiple output populations that are not encountered in healthy conditions, and may contribute to abnormalities in pain perception and reactivity.

The persistent expansion of nociceptive inputs to brain somatosensory circuits coupled with a loss of innocuous or discriminatory inputs likely plays a role in the generation of the affective and cognitive disturbances associated with chronic pain. Understanding how these different pathways interact and are distributed at brainstem, limbic, and cortical levels will clarify the impact of changes in the outflow from the spinal cord on the higher nociceptive circuit mechanisms and how this contributes to the complex sensory features of chronic pathological pain states.

### Distinct mechanisms for mechanical hypersensitivity dependent on pain state

Topical capsaicin, which induces a direct sensitization of the noxious heat transducer TRPV1 in those nociceptors that express it^8^, selectively potentiates responses to innocuous mechanical stimuli in the NOX population of ALT polymodal neurons that is normally activated only by high-intensity indentation stimuli. This may reflect an acute activity-dependent central sensitization such that subthreshold synaptic inputs from low threshold mechanoreceptors can now drive an output from these neurons. This result is consistent with a previous observation that responses to mechanical indentation in excitatory but not inhibitory lamina I neurons are potentiated by capsaicin injection in the mouse hindlimb^33^. The activation of normally high-threshold spinal projection output neurons by low-intensity stimuli closely aligns with the immediate hyperreactivity of mice to tactile stimuli after topical capsaicin treatment. In contrast, we did not observe an increase in low-threshold mechanical responsiveness after SNI, despite the induction of robust mechanical allodynia in injured mice. These data suggest that different pain states may differentially engage distinct ascending pathways to produce mechanical hypersensitivity^62,63^.

The absence of low-intensity mechanical recruitment in superficial ALT neurons in SNI is surprising given the evidence for spinoparabrachial neuron involvement in mechanical allodynia in other neuropathic pain models^23,64^. One possibility is that SNI may alter mechanosensory output through ascending pathways that do not directly involve the lamina I retPbN, Tacr1 or Gpr83 neuronal populations, such as deep lamina ALT neurons, the spinothalamic tract, or the dorsal column pathways. Changes in descending modulation of spinal circuits signals may also contribute to tactile allodynia following nerve injury, as may a temporal decorrelation of responses to touch in the dorsal horn^65^.

### Concluding remarks

As the primary conduit for pain-initiating and sustaining information to enter the brain, ALT neurons are an attractive target for understanding the mechanisms that drive and sustain pathological pain states which in turn may help the design of novel interventions to reduce these. We find that ALT neurons are functionally diverse, and that subtypes are differently but predictably impacted by distinct pain-triggering conditions and that the duration of these changes can be persistent. Targeting analgesic therapies to those specific cellular populations that transmit to the brain the abnormal sensory information that drives pathological pain is therefore a rational strategy. The approaches described here to identify and evaluate different ALT subpopulations should be readily extendable to a wide range of clinically relevant pain and itch models to identify sensory dysfunctions in specific neuronal spinal populations for a precise evaluation of the potential impact of specific genetic, pharmacological or surgical interventions on spinal projections to the brain to selectively prevent pathological sensory-input triggered brain disturbances in perception and mood.

## METHODS

### RESOURCE AVAILABILITY

#### Lead contact

Further information and requests for resources and reagents should be directed to the lead contact, Clifford Woolf.

#### Materials availability

This study did not generate new unique reagents.

#### Data availability

All data reported in this paper will be shared by the lead contact upon request.

#### Code availability

All computer aided design files used to generate 3D printed materials reported in this paper will be shared by the lead contact upon request, as will all code used for analysis and visualization of data.

### EXPERIMENTAL MODELS

All animal husbandry and all procedures involving mice were conducted in strict accordance with guidelines set forth by the Boston Children’s Hospital Institutional Animal Use and Care Committee. Mice expressing GCamP6f were generated by breeding of B6J.Cg-*Gt(ROSA)26Sortm95.1(CAG-GCaMP6f)Hze* mice (Ai95, JAX #28865)^37^ to either *Gpr83tm1.1(cre/ERT2)* mice (Gpr83^creER^, Jax #39021)^16^, *Tacr1tm1.1(cre/ERT2)* mice (Tac1^creER^, JAX #39022)^16^, or B6J.129S6(FVB)-*Slc17a6tm2(cre)* mice (Vglut2^cre^, Jax #28863)^66^. Adult mice of both sexes heterozygous for both GCaMP allele, and cre or creER alleles were used for imaging experiments, with initial age at implant ≥ 8 weeks. Induction of CreER translocation was performed at 5-8 weeks of age by three injections of 1mg tamoxifen in sunflower oil separated by 48 hours between treatments, with studies beginning at least 2 weeks after the last injection. Mice of both sexes heterozygous for the Ai95 allele were used for retrograde labeling of spinal projection neurons. Both male and female mice were included in all treatment groups for imaging experiments. Roughly equal numbers of C57BL6/J mice (Jax #000664) were used for comparison of Von Frey test with electronic plantar indentation assay, sex is specified in figure legend.

### METHOD DETAILS

#### Stereotaxic injection of Adeno-associated Virus

Bilateral injections of AAV2/retro-CAG-Cre-WPRE (6.21432E+13 gc/mL, Boston Children’s Hospital Viral Core) were targeted to parabrachial nuclei of Ai95 heterozygous mice for retrograde labeling of anterolateral tract neurons with GCaMP6f. Two 320nl injections of virus were made at coordinates (ML: 1.43mm AP: 5.25m) relative to bregma at dorsoventral coordinates of 3.5 and 3.7mm. Spinal window implantation was performed ≥ 4 weeks after virus injection.

#### Whole mount histology

Mice were perfused with ice cold 4% PFA. Spinal cords were dissected and placed into 4% PFA overnight at 4 °C then transferred to 1X PBS. Clearing solution was made by dissolving 20% (vol/vol) DMSO, 40% (vol/vol) TDE, 20% (wt/vol) sorbitol, and 6% (wt/vol) Tris base into ddH2)^67^. Spinal cords were removed from vertebral columns and cut into 1.5 cm sections. They were then placed into vials containing the clearing solution. Vials were placed on a shaker at room temperature shielded from light overnight, then stored at 4 °C. Cleared spinal cords were imaged on a Zeiss LSM 980 confocal microscope.

#### Spinal implant assembly

We incorporated elements of previously reported spinal fixation hardware^30,42^ to generate an assembly consisting of two lateral fixation bars mounted on snap-off delivery pins, and a baseplate that couples to the fixation bars to create and stabilize a window over the optically exposed spinal cord. These elements were first 3D printed using biocompatible resin (either Surgical Guide or Dental SG, Formlabs) using a Form 2 or Form 3B+ stereolithographic printer (Formlabs) at a 50-micron Z-resolution. Printed parts were washed in isopropanol for 10 minutes. Before further processing, stainless steel 000-120 hex nuts (McMaster-Carr) were press-fit into receptacles at each end of the fixation bars, and the delivery pins were attached to 6mm diameter stainless steel posts (Thorlabs) with 4-40 stainless steel set screws (McMaster-Carr). Mounting posts on baseplates were threaded with a 4-40 tap (McMaster-Carrr). All parts were then UV cured at 65 °C for 30 minutes, and then autoclaved in preparation for implantation. CAD files for all components are available from authors on request.

#### Spinal window implantation

Surgical procedures for window implantation were adapted from previously reported protocols^30,42^. Animals were weighed prior to surgery. Anesthesia was induced with 3.5% isoflurane in an induction chamber, eye ointment was applied, and dorsal hair clipped over surgical field, after which animals were transferred to a closed-loop heating pad maintaining rectally measured body temperature at 36-37 °C (FHC, Inc). Surgical plane anesthesia was maintained for the duration of the surgery with 1.5-2% isoflurane through a nose cone. Injections of 0.01ml/Kg of 5% dextrose in saline, 5mg/Kg meloxicam, and 0.2mg/Kg Dexamethosone were delivered intraperitoneally, and 100 microliters of 0.25% bupivicaine administered subcutaneously above the lumbar-thoracic vertebrae. The dorsum was then shaved closely with a safety razor and disinfected with three alternating administrations of povidone-iodine antiseptic (Betadine). At least 10 minutes after bupivacaine injection, the dorsal skin was incised and retracted from vertebral level T11 to L3. Tissue overlying targeted vertebral laminae typically (T13 and L1) was removed with forceps and the vertebral column freed by cutting lateral tendons at the point of attachment. Lateral fixation bars mounted on 6mm steel posts were then positioned using right-angle post adaptors (Thorlabs) to fix the vertebral column by compression. Blood flow through the spinal central vein visible between laminar segments was monitored to avoid excessive compressive force. Laminectomy was performed (typically at lamina T13 and L1) by careful thinning of the lateral edge of the laminae using a high-speed micro-drill (Roboz) driving 0.05mm tungsten carbide burrs (Fine Science Tools). Exposed spinal cord was washed with saline and kept moist for the remaining duration of surgery. The baseplate module of the 3D printed assembly was stably coupled to the fixation bars by screwing 3/32” length, 000-120 machine screws through slots in baseplate into the hex nuts at either end of the fixation bars. The laminectomy was then sealed with a 5mm diameter #0 glass coverslip (Warner Instruments) adhered to the laminectomy with Kwik-Sil silicone adhesive (World Precision Instruments). Any remaining gaps in assembly were sealed with dental acrylic (Lang Dental). Fixation bars were then snapped off from delivery pins and the surgical incision closed by adhering the margins to edges of implant with cyanoacrylate glue (Loctite). Animals were then removed from anesthesia and thermal support and monitored until they were locomoting normally. Implanted mice were singly housed in cages without food hoppers to avoid damage to the implant during healing.

For repair of windows exhibiting loss of optical clarity, mice were anesthetized with 1.5% isoflurane, eye ointment was applied, and thermal support was maintained, as for the implantation. The baseplate was immobilized by fixing mounting posts of baseplate to 6mm fixation posts, the previously implanted coverglass and elastomer were removed with forceps, and dura and overlying neoplastic tissues were removed with forceps and 26G needles and rinsed with sterile saline as needed for hemostasis. The window was then sealed with silicon elastomer and cover glass as for initial implantation.

#### Behavioral Assays

For all behavioral assays, mice were habituated in the test environment without an investigator present in the room for 1 hour on the day before the first test, and for 1 hour immediately prior to each test.

##### Von Frey assay

Habituation and testing were performed with mice in a small cage (7.5cm x 7.5cm x 15cm) on a mesh floor. The lateral surface of the paw was indented with nylon filaments with buckling forces selected according to an up-down protocol^68^. First stimulation was applied with an 0.6gcf buckling force filament, with filament stiffness iteratively increased on subsequent trials if non-responsive or reduced if responsive. Responses to mechanical indentation of the lateral paw were recorded as positive if the mouse lifted, shook, or licked the paw. Estimated 50% paw withdrawal methods were calculated from the response series as previously described^69,70^.

##### Electronic plantar indentation assay

Habituation and testing were performed in the same apparatus as for the Von Frey assay. For each mouse the lateral surface of each paw was indented 10 times with a flexible (buckling force ∼150mN) or rigid probe coupled to a digital force transducer (see Mechanical stimulation section for details). Each stimulus presentation was classified as withdrawal or non-withdrawal by the investigator. Withdrawal threshold for each mouse was defined as the median of peak force values recorded during withdrawal trials.

##### Hargreaves test

Sensitivity to heating was measured by assessing the latency of paw withdrawal to heating with a radiant heat source^71^. Animals were habituated in testing chambers on a 29 °C testing surface. During testing, a radiant heat source was directed to the targeted hindpaw until either paw withdrawal, or 30 seconds if no withdrawal was observed.

##### Topical capsaicin sensitization

For both Hargreaves tests and Von Frey tests of capsaicin induced sensitization, at completion of a 1-hour habituation period, mice were removed from the testing chamber, 0.1% capsaicin cream was applied topically to the dorsal and ventral surfaces of the hindpaw, and then wiped clean with a paper towel after 60 seconds. Mice were then placed back in the testing chamber for 15 minutes before beginning behavioral testing.

#### Spared nerve injury

Spared nerve ligation (SNI) surgery^28,46^ was performed under 2% isoflurane anesthesia. The tibial and common peroneal branches of the sciatic nerve were tightly ligated with a 5.0 silk suture and transected with a 3-5mm section removed distally from each nerve. The sural nerve was left intact. The surgical incision was closed with sutures and mice recovered on heated pads before being returned to their home cage.

#### Calcium imaging

##### Anesthetized dorsal horn calcium imaging

Animals were anesthetized with 3% isoflurane in an induction chamber and subsequently maintained at 1% isoflurane via a nose cone. Ophthalmic ointment was applied, and the animal was placed on an imaging stage with closed-loop temperature control via rectal thermometer and resistive heating pad (FHC, Inc.), maintaining body temperature at 36-37 °C. For each animal’s first imaging session, a circular imaging adapter printed in black resin (Formlabs) was fixed to the implanted baseplate mounting posts with 4-40 set screws (McMaster-Carr). The imaging adaptor was immobilized on the imaging stage via 6mm steel posts and placed under an SFMGFP2 upright microscope (Thorlabs). The targeted hindpaw was prepared for stimulation as described below, either placed in a thermal stimulation chamber or bonded temporarily to a mechanical stimulation platform.

The imaging field was illuminated with a camera-exposure triggered 470nm LED light source using a 10X magnification, 0.3 numerical aperture, 10mm working distance air objective (Olympus). Excitation and emission light for GCaMP imaging were filtered with a standard GFP filter cube (Excitation: 469 ± 17.5nm, Emission: 525 ± 19.5nm, Dichroic reflection band 452-490, transmission band 505-800). Fluorescence images were recorded with a CMOS camera (1920 x 1080 pixels, 5.04 micron pixel width and height, Thorlabs) at 10Hz with 99ms exposure time. Images were time-stamped for synchronization with other experimental data streams.

For repeated imaging of specified areas in chronic imaging experiments, imaging fields were located by visualization of vasculature imaged under reflected light. The orientation of the imaging window was rotated around mediolateral and rostrocaudal axes to level the imaging field within the focal plane of the objective lens.

##### Awake dorsal horn calcium imaging

Mice were habituated to tail-held walking on a mesh treadmill under imaging microscope for 10 minutes 1 day before first imaging session, in a darkened room. For imaging, awake mice were spine fixed on a treadmill by coupling the same imaging adaptor used for anesthetized imaging to transverse fixation posts. Imaging was performed in the dark with targeted illumination of the hind paws with white light. Imaging parameters were the same as for anesthetized mice, with the exception of stimulation protocols as described below. For direct comparison of awake and anesthetized responses within the same imaging field, awake imaging was performed prior to anesthetized imaging.

##### DRG calcium imaging

Male and female mice (8 -12 weeks) carrying both Ai95 and Vglut2^cre^ alleles were prepared for imaging of dorsal root ganglia. Anesthesia was induced with 2.5% isoflurane and maintained with 1.5% isoflurane delivered via a nose cone. Mice were placed on a heated pad to maintain the body temperature at 37 °C, and ophthalmic ointment was applied to the eyes throughout the surgery and imaging. The back was shaved, and a rectangular incision was made between the lumbar enlargement and pelvic bone with the skin folded away. Dorsal laminectomy was performed in four steps. First, a 1.5-cm-long midline incision caudal to the lumbar enlargement area was made to create space for clamping. Second, a 3-4 mm incision was made approximately 1 cm rostral to the pelvic bone to expose the spine where the L4 DRG is situated. Third, muscles and connective tissues were cleaned with scissors and forceps. Fourth, the L4 transverse process was removed using rongeurs and fine forceps to expose the DRG. To minimize movements caused by breathing, a custom spinal clamp was used to secure the spine over the vertebrae 5mm rostral to the L4 DRG. Illumination was provided with a collimated 470nm light-emitting diode through a 10x, 0.3 numerical aperture air objective. The clamp was adjusted to ensure the DRG surface was as level as possible, visualizing the maximum surface area in the focal plane (typically covering ∼80% of the entire DRG). Images were acquired as for anesthetized spinal imaging (10 Hz with a 99 ms exposure time). Isoflurane concentration was reduced to 1% during the imaging process. For each animal, thermal and mechanical stimulation, and capsaicin application were performed following the same procedures as described for anesthetized spinal imaging.

#### Somatosensory stimulation

##### Thermal stimulation

Methods for generating thermal stimuli were adapted from previously reported methods^31,55^. A pressurized reservoir containing distilled water was connected to 2 digitally controlled Peltier liquid cooling-heating units (TE Technology), one set permanently to 35 °C and the other modulated from -1 °C to 51 °C during stimulus delivery. For anesthetized imaging, water flow from the temperature control units was directed to a stimulation chamber in which the targeted hindpaw was submerged, with flow regulated by solenoid pinch valves under microprocessor control (Arduino). For awake imaging, stimulation flow was directed to the targeted hindpaw from below the mesh treadmilll. For each stimulus temperature, the paw was stimulated with 16 seconds of 35 °C water interspersed with 8 seconds flow from the modulated temperature source. Temperature of water at the paw was monitored with a radiometric FLIR camera unit (Lepton 3.5, Sparkfun), with each thermal image frame time-stamped for synchronization with GCaMP imaging data.

##### Mechanical stimulation

Quantitative indentation was applied manually using a custom instrument consisting of an100g capacity load cell (Sparkfun) coupled at one end to a 3D printed ergonomic handle and at the other to a magnetic probe adaptor, allowing exchange of indentation probes with varying properties. For low to moderate intensity stimulation, the device was coupled to 480 micron diameter nylon filament with buckling forces ranging from 20 to 150 millinewtons. For intense stimulation the device was coupled to a 630 micron diameter stainless steel rod. Force output from the load cell was digitized and visualized online to deliver controlled indentation stimuli. Stimulation was simultaneously recorded at 25-30fps with a digital camera (Spinel). Force measurements and video output were time-stamped for synchronization with GCaMP imaging data. For anesthetized imaging experiments, the hindpaw was fixed in place during mechanical stimulation with latex adhesive. Stimuli were applied for 1-5 seconds to at least 5 locations within each sural, tibial, and saphenous receptive fields of the glabrous paw skin, or sural, peroneal, and saphenous fields of hairy skin. For awake experiments indentation was applied from above to stimulate hairy paw skin or from below through mesh treadmill to stimulate glabrous skin.

#### Calcium imaging data analysis

##### Pre-processing

Image series were median filtered to correct bad pixels. A low-pass filtered image series was generated and subtracted from the original series to correct for global changes in illuminance across the imaging area and global luminance over time (e.g. photobleaching). Planar motion within imaging sessions was corrected by applying rigid image registration using the StackReg library^72^. Candidate regions corresponding to active cells were first identified by adaptive thresholding of heat maps generated by correlation of pixels to their nearest neighbor. These masks were used as input for constrained non-negative matrix factorization^73^ to generate refined regions of interest. From these, GCaMP fluorescence traces were calculated as mean value of pixels in the spatial footprint over imaging time-series, and modified z-scores calculated by normalization to median absolute deviation of the trace, multiplied by k = 1.4826 for comparability to standard deviation. Significant calcium transients were identified as trace values above a z-score of 3 for a duration of at least 200ms.

##### Multi-session image alignment

Following within-session image registration and segmentation, image time series and segmented regions of interest were aligned across sessions by rigid-body registration of median frame images using StackReg. Multi-session series were cropped to include only areas visible in all recordings. The multi-session alignment function of the CaImAn library was used to identify neurons conserved across sessions^74^.

##### Thermosensory correlation visualization

Correlation maps were constructed by regressing GCaMP image series pixel intensities to temporally aligned stimulus temperature values and assigning the absolute value of the correlation coefficient to the map location of each pixel^75^. Overlaid images showing multiple stimulus correlation maps were generated by additive blending of component maps.

##### Clustering of thermosensory responses

A response matrix was created by aligning GCaMP responses of retPbN cells to first presentation of stimuli at 8 °C, 23 °C, 39 °C, 42 °C, 45 °C, 47 °C and 50 °C (±1 °C). Silhouette analysis identified a peak at k=3 clusters, therefore k-means clustering was performed on the response matrix with k=3, and cluster centers were calculated and aligned to stimuli used to generate the original response matrix. Principal component analysis was performed using the SciPy stats module, with the first 10 components reported.

##### Thermosensory tuning polar plots

For each presentation of a thermal spectrum to a cell, a response vector was created from the median response to repeated presentations of nominal 10 C, 42, and 50 C stimuli. This vector was projected along orthogonal axes and then transformed to polar coordinates to calculate thermal tuning angle.

##### Mechanosensory response analysis

Traces of recorded probe force were segmented into individual stimuli by thresholding and identification of peaks. Calcium transients (z-score value above 3 for ≥ 200ms) were assigned to mechanical stimuli during which they were initiated. For receptive field mapping, each mechanical stimulus peak was aligned to probe position recorded synchronously in video and manually located on a map of the paw. The center of the receptive field was calculated as the centroid of the convex hull surrounding locations at which responses above threshold were evoked and color-coded according to a color gradient superimposed on the paw map.

### QUANTIFICATION AND STATISTICAL ANALYSIS

Statistics were performed using SciPy’s stats module (Python 3.9) or GraphPad Prism and used to make comparisons between groups. Group sizes and units are specified in figure legends. Normality assumptions were tested by Shapiro’s method before applying parametric tests. In all figures, significance is defined as *p= < 0.05, **p= < 0.01, ***p<0.001. Error bars display 95% confidence intervals unless otherwise noted.

## SUPPLEMENTAL FIGURE LEGENDS

**Supplemental Figure 1.**
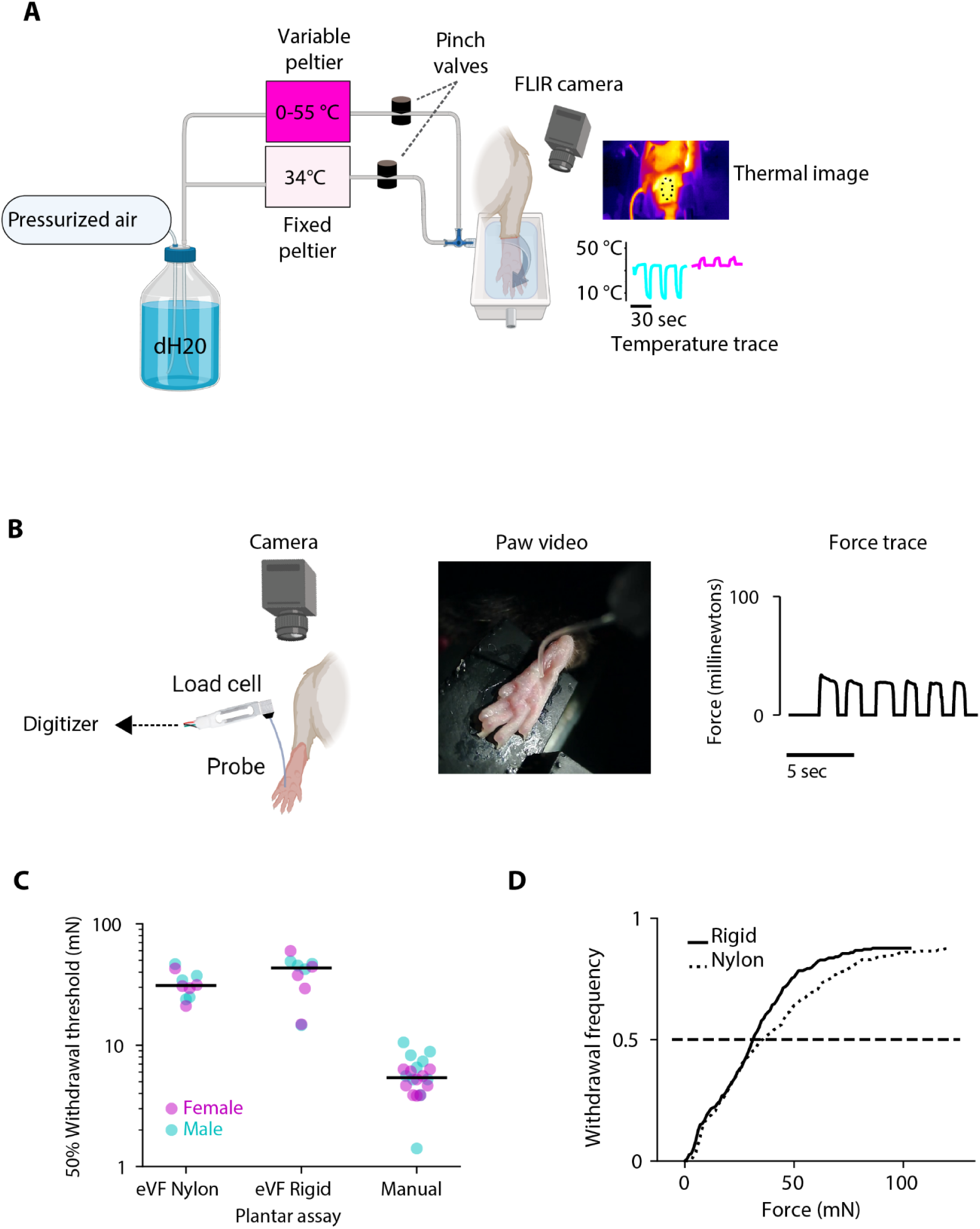
Quantitative somatosensory stimulation of the hindpaw. A. Illustration of thermal stimulation. Water is pumped through two Peltier-controlled chiller/heater units and is directed through tubing to a hindpaw placed in a stimulation chamber (left). Water flow is controlled by solenoid valves to alternately activate a constant neutral temperature source (35 °C) or variable source (0-55 °C). A radiometric thermal camera records temperature at the paw (right, thermal traces extracted from the circled region are shown below). B. Illustration of mechanical stimulation protocol. Glabrous hindpaw is indented with a probe coupled to a shear force transducer. A digital camera records stimulus location in synchrony with force measurement. C. Force intensity eliciting a 50% likelihood of paw withdrawal on indentation of the hindpaw with the flexible and rigid probes used in the imaging experiments and assessed by standard manual Von Frey assay (up/down method). Data represent the same cohorts of 5 female and 5 male animals assessed with each of the three methods. D. Paw withdrawal probability versus indentation force for naive mice stimulated with rigid steel or flexible nylon probes used for mechanical stimulation in imaging experiments.

**Supplemental Figure 2.**
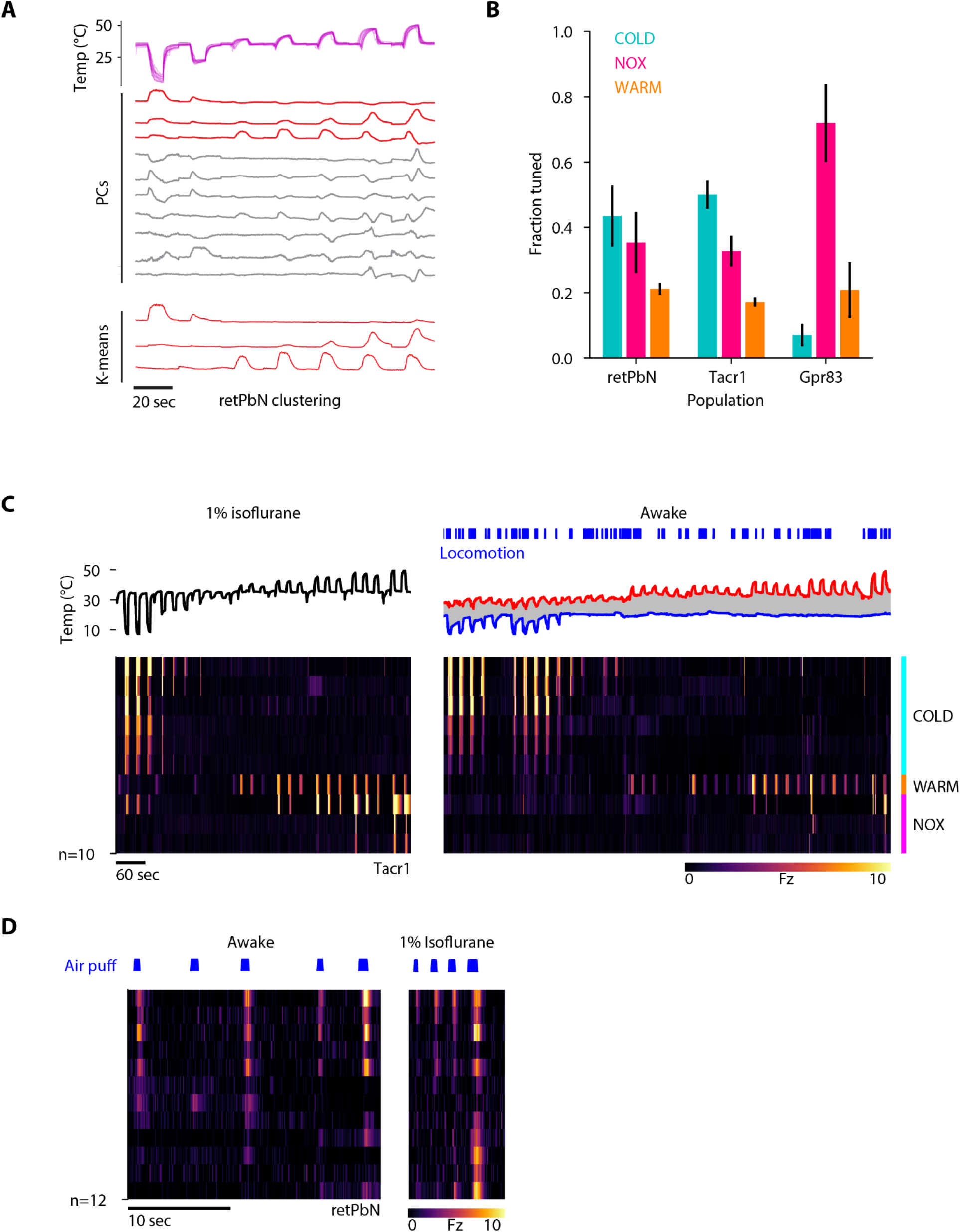
Clustering of thermosensory response types. A. Identification of prototypical thermosensory response types among retPbN neurons using principal component analysis (top) and K-means clustering with K=3 (below), applied to a stimulus-aligned GCaMP response matrix (see Figure 2B). Stimuli presented to generate the response matrix are superimposed above. First 10 principal components and 3 K-means centers are shown, with those used to definethe three thermosensory classes highlighted in red. B. Distribution of thermosensory response types in retPbN, Gpr83 and Tacr1 neurons. Thermally responsive cells are classified according to tuning angle illustrated in Fig 2F-G. GCaMP activity recorded from N=11 Tacr1, 5 GPr83 and 9 retPbN mice. C. Rasters of GCaMP fluorescence of the same set of Tacr1 neurons elicited by cooling and heating of the ipsilateral hindpaw recorded under 1% isoflurane anesthesia (left) or when awake on a treadmill (right). In both cases stimuli were applied with a constant pressure, temperature modulated stream of water directed onto the hindpaw. The temperature trace for the awake condition plots minimum and maximum temperatures measured in the region occupied by the ipsilateral hindpaw. Locomotion during imaging is plotted above the awake raster. D. Raster of retPbN neuron GCaMP responses to stimulation of the ipsilateral hindpaw with cold compressed air in the same set of neurons recorded either in an awake, spine-fixed animal (left), or under 1% isoflurane anesthesia (right).

**Supplemental Figure 3.**
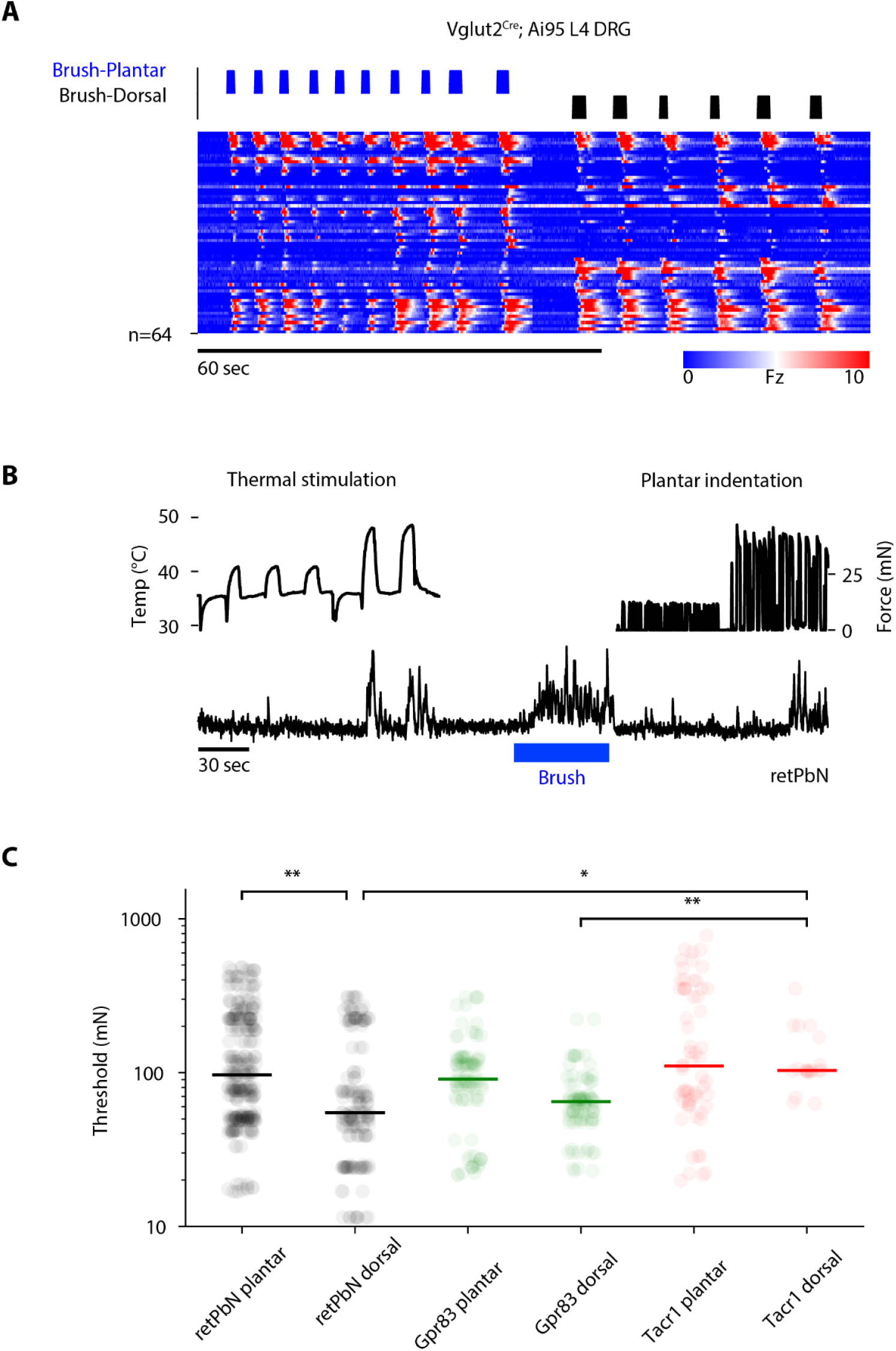
Sensitivity of DRG and lamina I neurons to mechanical stimulation of hindpaws. A. Response raster of primary sensory neurons in L4 DRG of Vglut2^Cre^;Ai95 mice in response to brushing the plantar and dorsal ipsilateral hindpaw. B. GCaMP fluorescence trace of a mechanically responsive retPbN neuron in response to heating, brushing, and low-to moderate-intensity indentation of the hindpaw. C. Response thresholds of dorsal horn neurons to indentation of plantar and dorsal surfaces of the hindpaw. Bars represent the median of responses for each group. Responses were obtained from 6 Tacr1, 5 Gpr83, and 7 retPbN mice. Comparisons made by Krustal-Wallis test followed by Dunn’s multiple comparisons test.

**Supplemental Figure 4.**
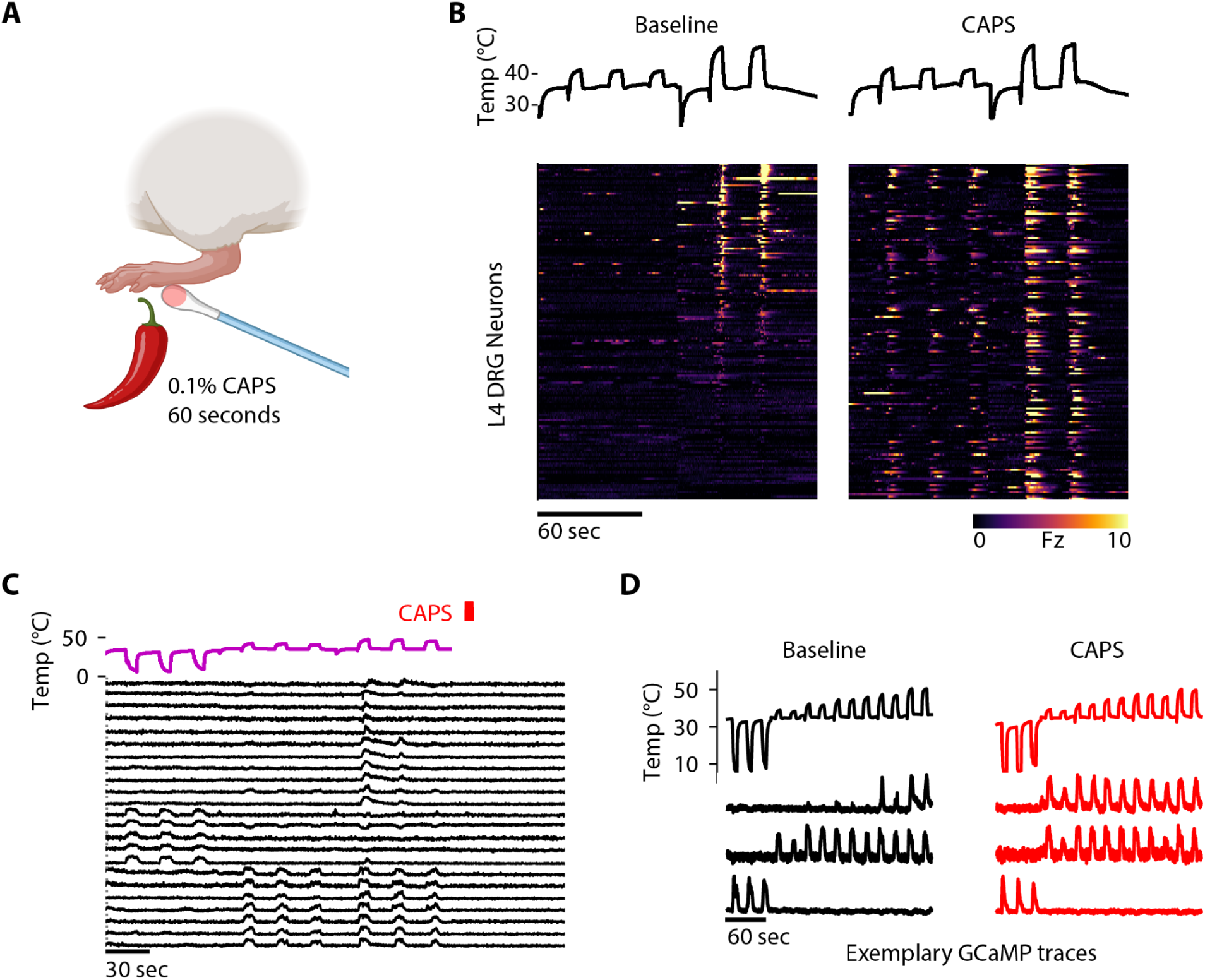
Characterization of topical capsaicin sensitization model. A. Topical capsaicin sensitization model. Capsaicin ointment (0.1%) was applied to the skin of one hindpaw with a cotton swab and removed after 60 seconds. B. Rasters of GCaMP activity evoked in Vglut^cre^ L4 dorsal root ganglion neurons by heating of the hindpaw, before and directly after topical capsaicin treatment. C. Examples of GCaMP fluorescence traces from three retPbN neurons corresponding to each of the thermosensory classes (top NOX; middle WARM and bottom COLD) in response to a full range of thermal stimuli shown above the traces, before and after topical capsaicin treatment.

**Supplemental Figure 5.**
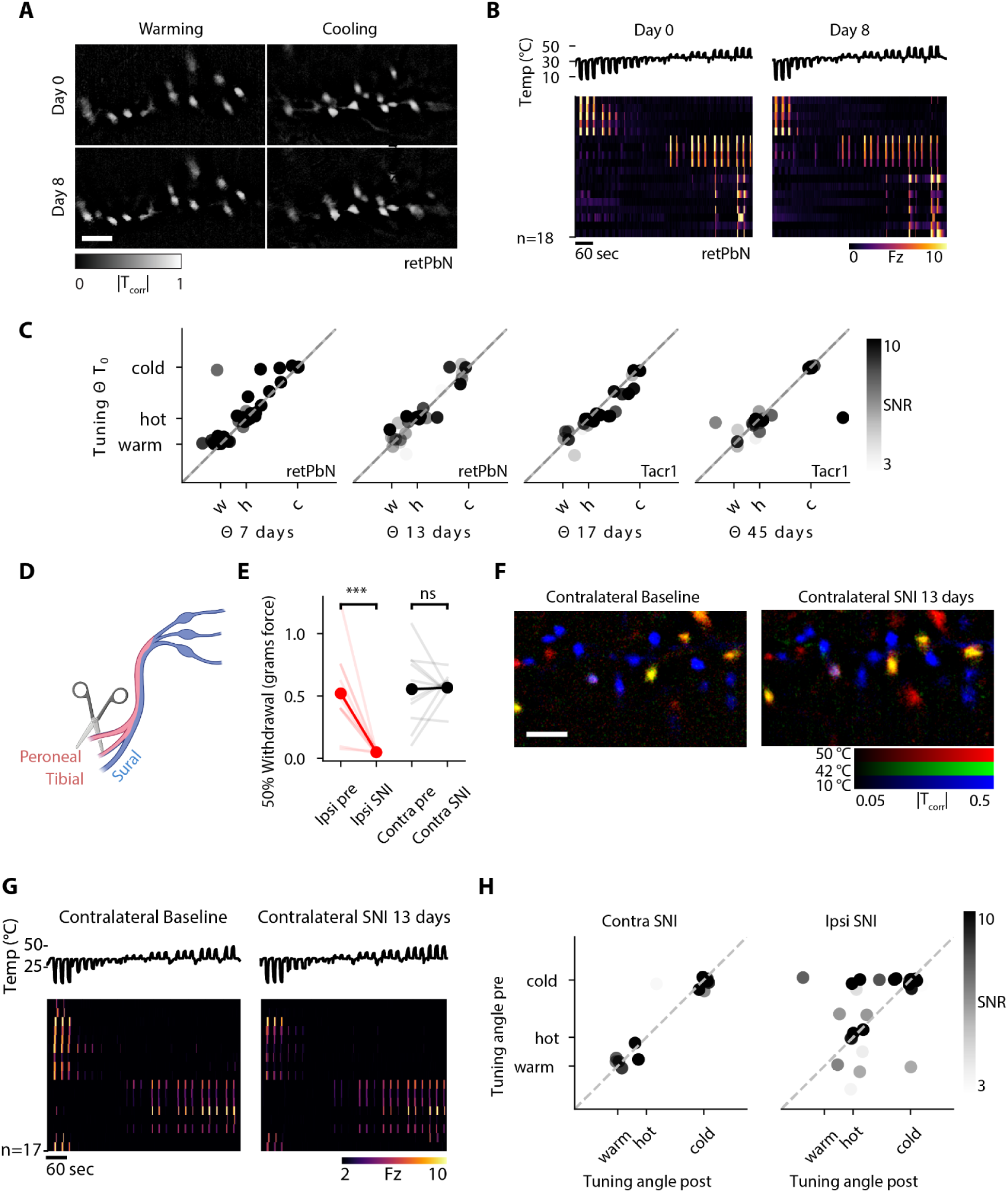
Stability of spinal sensory tuning in the absence of injury. A. Spatial maps of correlation between retPbN GCaMP fluorescence responses to cooling or warming stimuli recorded within the same imaging field 8 days apart. B. Rasters of GCaMP fluorescence in response to cooling and heating stimuli recorded from the same retPbN neurons, recorded 8 days apart. C. Scatter plot of thermosensory tuning angles, (as determined for Fig 2F-G) of neurons recorded at different time intervals. Mean signal to noise ratio of responses (SNR) is displayed for each neuron. D. Diagram of spared sural nerve injury (SNI). The peroneal and tibial branches of the sciatic nerve are transected and ligated, leaving intact the sural nerve, which innervates the lateral surface of the hindpaw. E. Paired plot of mechanical forces eliciting a 50% paw withdrawal probability for mice used in SNI imaging studies ipsilateral (red) and contralateral (black) to the nerve injury, measured prior to and 4 weeks after injury. Paired t-test, N=11 mice. F. Maps of correlation between GCaMP fluorescence and thermal stimuli overlaid for an imaging field before and 13 days after SNI contralateral to the imaging site. Scale bar is 200 microns. G. Raster of GCaMP responses for cells from the imaging field shown in panel F. H. Scatter plot of thermosensory tuning angle of neurons recorded before and 13-14 days after SNI ipsilateral and contralateral to the imaging site. Mean signal to noise ratio of responses (SNR) is displayed for each neuron.

**Supplemental Figure 6.**
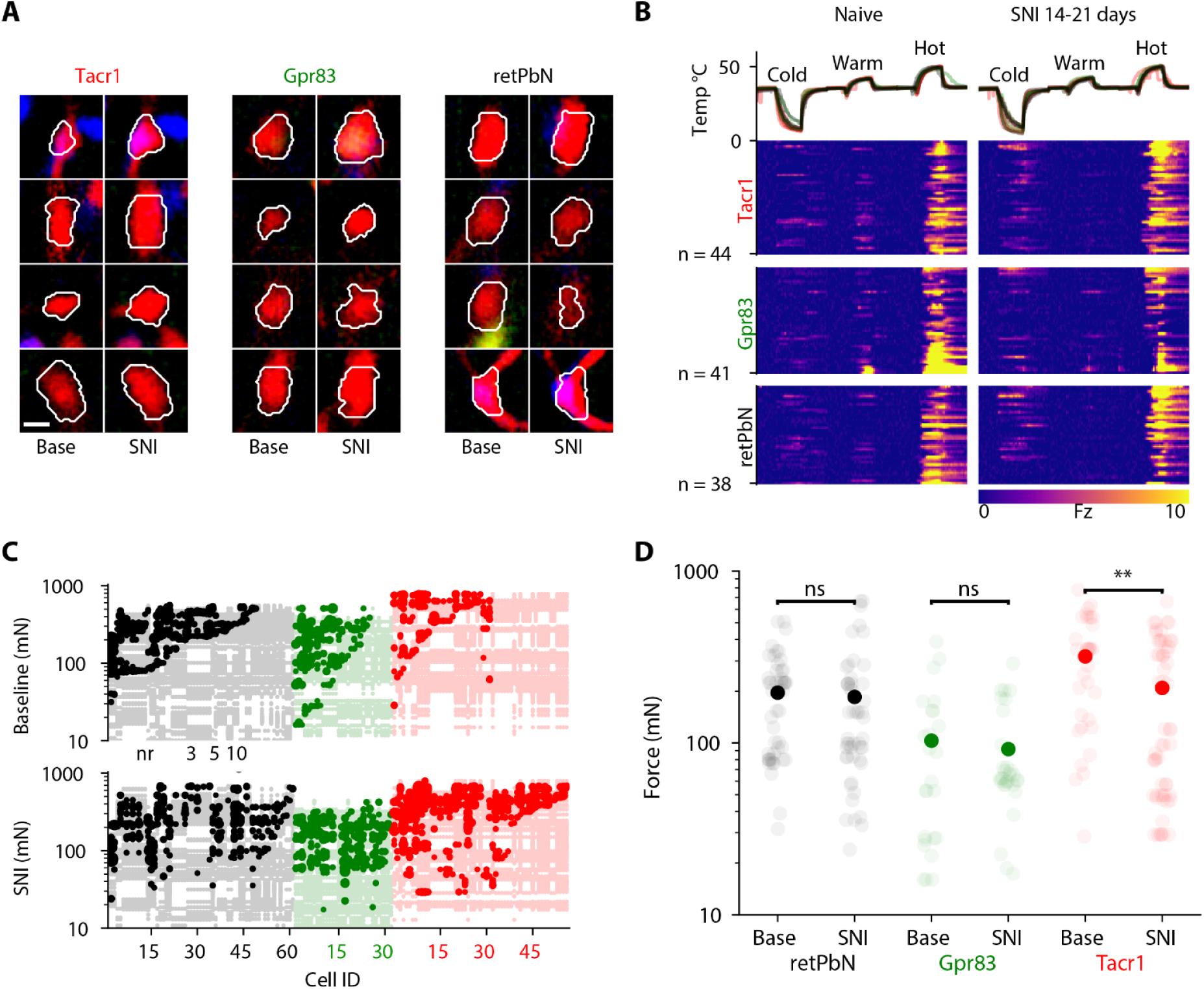
Sensory tuning of basal NOX neurons is conserved after SNI. A. Maps of the correlation between GCaMP fluorescence and thermal stimuli for NOX neurons in the Tacr1, Gpr83, and retPbN populations. Paired images show imaging locations aligned between sessions recorded before and 14-21 days after ipsilateral SNI. Contours of the regions used to extract calcium signals are overlaid for each image. Scale bar is 25 microns. B .Rasters of GCaMP activity recorded from Tacr1, Gpr83, and retPbN NOX neurons aligned to cooling, warming, and heating thermal stimuli, before (left) and 14 days after (right) induction of SNI ipsilateral to imaging field. GCaMP activity recorded from N=4 Tacr1, 3 GPr83 and 4 retPbN mice. C. Plot of all recorded responses to indentation before and 2-3 weeks after SNI. Individual cells are plotted on x axis, each point corresponds to a single stimulation, marker size corresponds to response amplitude for responses above threshold (fluorescence z-score ≥ 3 for 200ms), responses below threshold are identified by light shading. GCaMP activity recorded from N=3 Tacr1, 3 GPr83, and 3 retPbN mice. D. Minimum indentation force evoking GCaMP response in retPbN, Gpr83 and Tacr1 indentation-sensitive neurons recorded at baseline and 2-3 weeks after SNI. Mean values shown in solid color, comparisons made by Mann-Whitney U test.

**Supplemental Movie 1**

GCaMP fluorescence responses visualized in a retPbN mouse 14 days after spinal window implantation under 1% isoflurane anesthesia, in response to thermal stimulation of the ipsilateral hindpaw (overlaid). Imaging field is 1032 x 580 microns.

**Supplemental Movie 1**

GCaMP fluorescence responses from the same imaging session as in Sup. Movie 1, in response to quantitative indentation of the ipsilateral hindpaw (overlaid).

## ACKNOWLEDGEMENTS

We thank Bruna Lenfers Turnes for assistance with the spared nerve injury surgeries, and Nick Andrews for help with stereotactic injections. We thank Ofer Mazor and Pavel Gorelik from the HMS Research Instrumentation Core for technical support, and thank Seungwon Choi, David Ginty and members of the Ginty and Woolf labs for helpful comments on the manuscript. Illustrations were created with Biorender.com. We thank the Boston Children’s Hospital IDDRC Cellular Imaging Core, funded by NIH P50 HD105351, and the Boston Children’s Hospital Viral Core,, funded by NEI P30 grant 5P30EY012196.This work was supported by NIH grants K99DE028360 from the NICDR (DAY), AT011447 from NCCI (CJW) and R35NS105076 from NINDS (CJW).

## AUTHOR CONTRIBUTIONS

DAY and CJW conceived the study. DAY developed a chronic *in vivo* spinal imaging platform and performed imaging experiments with support from CG. XZ performed dorsal root ganglion imaging experiments. Histological analysis was performed by NMB and DAY. Behavioral experiments were performed by CK, CG, and NMB. DAY analyzed all data with assistance from CG. DAY and CJW wrote the paper with input from all authors.

